# An autophagy program that promotes T cell egress from the lymph node controls responses to immune checkpoint blockade

**DOI:** 10.1101/2023.07.17.549282

**Authors:** Diede Houbaert, Apostolos Panagiotis Nikolakopoulos, Odeta Meçe, Kathryn Jacobs, Jana Roels, Gautam Shankar, Madhur Agrawal, Sanket More, Maarten Ganne, Kristine Rillaerts, Louis Boon, Magdalena Swoboda, Max Nobis, Larissa Mourao, Francesca Bosisio, Niels Vandamme, Gabriele Bergers, Colinda LGJ Scheele, Patrizia Agostinis

**Affiliations:** Cell Death Research and Therapy Group, Department of Cellular and Molecular Medicine, KU Leuven, Herestraat 49, 3000 Leuven, Belgium; VIB Center for Cancer Biology Research, 3000 Leuven, Belgium; Laboratory of Intravital Microscopy and Dynamics of Tumor Progression, Department of Oncology, KU Leuven, 3000 Leuven, Belgium; Laboratory of Tumor Microenvironment and Therapeutic Resistance Center for Cancer Biology, Department of Oncology, KU Leuven, 3000 Leuven, Belgium; VIB Single cell core, Ghent/Leuven, Belgium; Laboratory of Translational Cell and Tissue Research, Department of Pathology, KU Leuven and UZ Leuven, Belgium; JJP Biologics, Warsaw, Poland; Intravital Imaging Expertise Center, VIB Center for Cancer Biology, VIB, Leuven, Belgium

**Keywords:** autophagy, lymphatic endothelial cells, lymph node, T cell trafficking, cancer, immunotherapy

## Abstract

Lymphatic endothelial cells (LECs) lining the lymphatic vessels of the lymph node (LN) parenchyma orchestrate leukocyte trafficking and peripheral T cell dynamics. T cell responses to immunotherapy largely rely on peripheral T cell recruitment in tumors. Yet, a systematic and molecular understanding of how LECs within the LNs control T cell dynamics under steady state and tumor-bearing conditions is lacking. Using intravital and high-resolution imaging combined with immune phenotyping, we show that LEC-specific deletion of the essential autophagy gene *Atg5* alters intranodal positioning of lymphocytes and accrues their persistence in the LNs, by increasing the availability of the main egress signal S1P. Single-cell RNA-sequencing of tumor-draining LNs from WT and ATG5^LEC-KO^ mice unveils that loss of ATG5 remodels niche-specific LEC phenotypes, involved in molecular pathways regulating lymphocyte trafficking and LEC-T cell interactions. Functionally, loss of LEC-autophagy prevents recruitment of tumor-infiltrating T cells and NK cells and abrogates tumor regression in response to anti-PD-1 or anti-CTLA4-based immunotherapy. Thus, a unique LEC-autophagy program boosts immune-checkpoint responses by guiding systemic T cell dynamics.

**Graphical Abstract:** 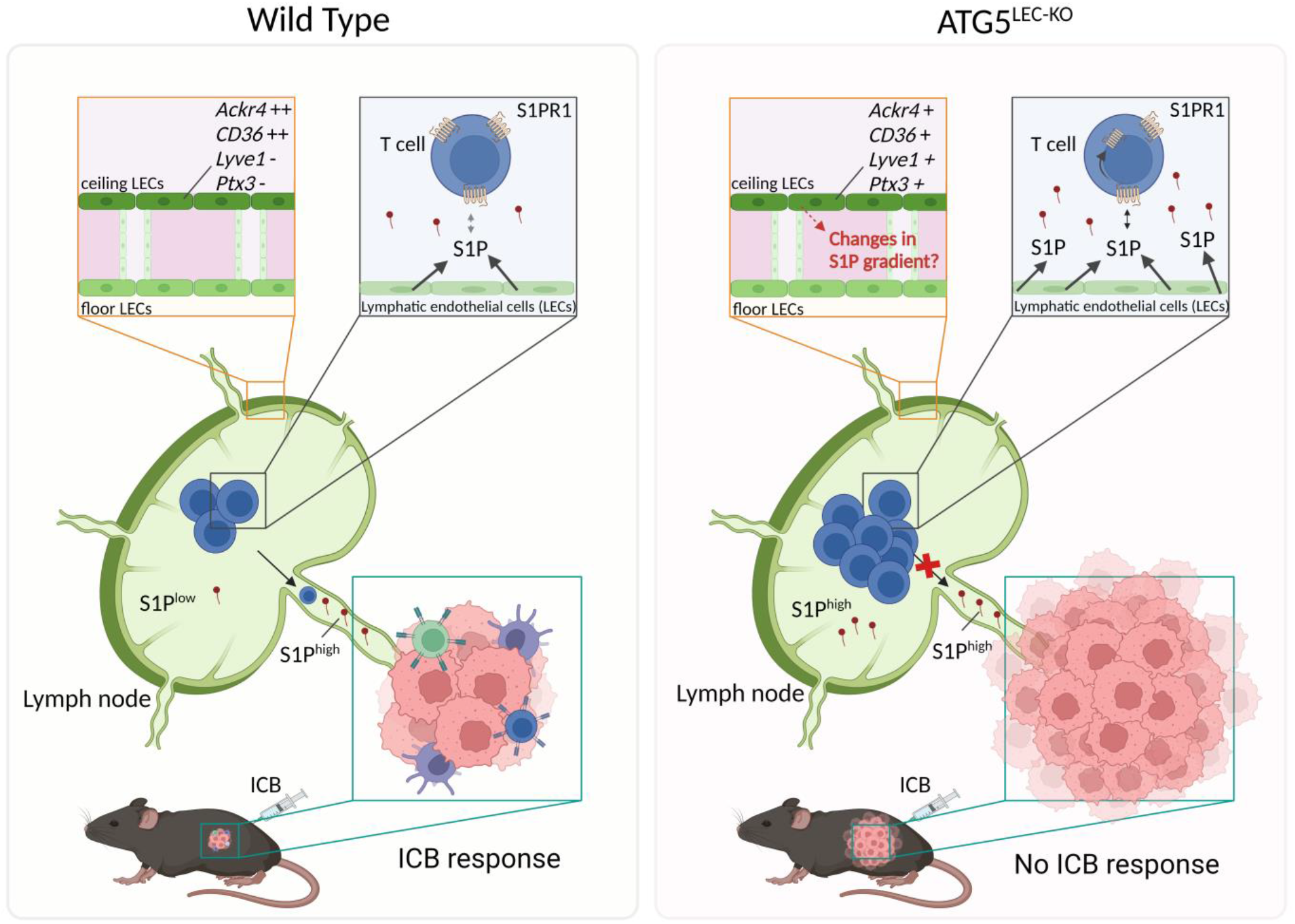

## INTRODUCTION

Lymph nodes (LNs) are immunological centers that orchestrate antigen acquisition and immune challenge, enabling the generation of appropriate adaptive immune responses against foreign invaders or cancer cells expressing abnormal self-/neo-antigens. These vital immunological functions require tight spatiotemporal coordination, involving multidirectional interactions between large pools of lymphocytes, which constantly recirculate from the blood through LNs in search for antigens and back, and specialized cellular components of the LN parenchyma. The LN parenchyma’s discrete anatomic and functional niches and the main regulators of the entry and egress of lymphocytes into and out of the LN have been well-characterized (1–4). After entering the LNs from the bloodstream, through high endothelial venules (HEVs) or via afferent lymphatics (5,6), major HEV- and stromal cell-derived lymphoid chemokines CCL21, CCL19 (for T cells and dendritic cells (DCs)) and CXCL13 (for B cells) guide intranodal positioning of naïve lymphocytes, which traffic CCR7+ T lymphocytes (and DCs) to the paracortical T cell areas and CXCR5+ B cells follicles to the cortex (7). Naïve lymphocytes that did not encounter their target antigens and activated T cells leave the LNs via medullary sinuses through efferent lymphatic vessels (8). The permanence of lymphocytes within the LN parenchyma is regulated by the balance of CCR7-ligands CCL21 and CCL19, which function as the main retention signals, and sphingosine-1-phosphate (S1P), the main egress-promoting signal secreted through the S1P transporter Spinster homolog-2 (SPNS2) by lymphatic endothelial cells (LECs). S1P concentration is usually high in the lymph fluid and plasma but is kept low in the LNs due to the action of S1P degrading enzymes (4,9), thereby creating a gradient that guides the exit of lymphocytes from the LNs back into circulation (8). S1P is sensed by a class of seven-transmembrane G-protein coupled S1P receptors (S1PRs) on lymphocytes, which upon binding to S1P become internalized, leading to their egress from the LN. Congruently, adoptively transferred *S1Pr1*^-/-^ T and B cells can enter the LN but cannot exit into the lymph (10). Likewise, alterations of the S1P gradient, either by deletion of S1P- generating sphingosine kinases in LECs (11) or by inhibition of S1P lyase, which degrades S1P (9,12), result in lymphocytes retention in the LN and consequent lymphopenia, highlighting the relevance of LEC-generated S1P as main egress-signal during homeostasis. Furthermore, the mechanism of action of the potent immunosuppressive drug FTY720 (fingolimod) relies on the downregulation of S1PR1 on lymphocytes and their segregation in the LNs, which improves autoimmune T and B-cell responses (13). While mechanisms governing T cell recruitment and egress from peripheral blood into LNs have been well documented (14), molecular processes regulating S1P production by LECs have not been characterized to the same extent. Also, it remains largely unknown how the unprecedented LEC heterogeneity that emerged from recent single-cell RNA sequencing (scRNAseq) analyses of murine and human LNs (15–17), impacts T cell dynamics and LN immune responses, both in resting and pathological conditions or during therapeutic interventions.

In cancer, tumor-draining LNs (TdLNs) are critical anatomic sites where naïve CD8 T cells traffic to be primed by DCs, prior to their migration through the blood as effector CD8 T cells, to infiltrate the tumor parenchyma (17). Moreover, recent reports have indicated that during cancer immunotherapy, TdLNs are the first site of T cell priming generating tumor-reactive T cells, which then traffic to the tumor parenchyma to endorse anti-tumor immunity, leading to tumor regression (18–21). Congruently, in melanoma patients, T cell proliferation in peripheral blood, rather than of tumor-infiltrating T cells (TILs), is associated with greater responses to anti-PD-1 blockade (22). Altogether, these studies suggest that mechanisms regulating peripheral T cell dynamics critically shape immunotherapy responses. Thus, insights into the mechanisms that control lymphocyte egress from the LNs are key to our ability to control systemic anti-cancer immune responses.

Recent studies from different laboratories including ours uncovered that macroautophagy (from here on referred to as autophagy) functions as a primary process controlling lymphatic vessel homeostasis and responses of LECs to inflammatory cues (23–25). Autophagy is the major lysosomal pathway for the degradation and recycling of cytoplasmic components, with emerging cell-specific functions that are modulated in response to intrinsic and extrinsic cues or stressors (24,26). We recently reported that the genetic blockade of autophagy in LECs remodels lipid metabolic pathways and prevents inflammation-driven corneal lymphangiogenesis (25). In contrast, in a model of collagen-induced arthritis (CIA), the genetic deletion of ATG5 in LECs aggravated CIA-driven inflammation, by reducing the proliferation of regulatory T cells in the LNs from arthritic mice and by stimulating pathogenic Th17 cell egress from the LNs (27). While these observations support the highly context-specific function of autophagy, it has not been studied systematically whether LEC-autophagy regulates LN homeostasis and anti-tumor immune responses.

Using intravital imaging and immune profiling, we unveiled that genetic deletion of ATG5 in LECs traps lymphocytes in secondary lymphoid organs (SLOs), both under steady-state conditions and in tumor-bearing mice, by altering the gradient of the egress signal S1P within the LN parenchyma. In line with this, by combining spatial analysis and single-cell RNA sequencing (scRNAseq) of TdLNs, we uncovered that loss of LEC-autophagy remodels LN LEC subtypes and lymphocyte positioning within the LN parenchyma. Deletion of ATG5 in LECs abrogates tumor regression in response to the main immune checkpoint blockers (ICB), anti-PD-1 and anti-CTLA4, by preventing the migration of newly ICB-responsive T cell and NK cells to the tumor. Thus, LEC-autophagy is required to maintain the recirculation of lymphocytes through the LNs, which is of paramount relevance for the systemic anti-tumor immune responses elicited by ICB.

## RESULTS

### Genetic deletion of ATG5 in LECs traps lymphocytes in secondary lymphoid organs

To assess the role of LEC-autophagy in the LNs, we conditionally knocked out ATG5 in the lymphatic endothelial compartment (ATG5^LEC-KO^) by crossing *Atg5*^fl/fl^ mice with *Prox1*-*cre^ERT2^*mice expressing a tamoxifen-inducible Cre recombinase, as reported in our previous study (25). Staining for LYVE1+ lymphatic vessels of the inguinal LN indicated that compared with wild-type (WT) mice, ATG5^LEC-KO^ mice displayed a lack of ATG5 signal and reduced autophagosomal-bound LC3 immunoreactivity, pointing to impaired autophagy flux (Supplementary Fig. 1A,B). Lymphatic vessel density in the LNs was not altered by genetic deletion of ATG5 in LECs under steady-state conditions (Fig. 1A). This suggests that in the absence of inflammatory signals driving remodeling of lymphatic vessels and lymphangiogenesis in LNs, impairing autophagy in LECs does not lead to major changes in the lymphatic vessel network. Immunostaining for the pan-T cell marker CD3 in thin sections (Supplementary Fig. 1C) and in intact LNs of WT and ATG5^LEC-KO^ mice crossed with R26-mTmG mice, expressing membrane-localized tdTomato fluorescence in all tissues/cells (Fig. 1B, Video 1,2), revealed an increase in cellularity (total area of CD3^+^ cells) in LNs of ATG5^LEC-KO^ mice compared with WT mice. Congruently, the number of both naïve (CD62^high^CD44^low^) CD4 T and CD8 T cells expanded in the LNs of LEC-ATG5 deficient mice (Fig. 1C). A similar phenotype was observed in the spleen of these mice (Fig. 1D), indicating that impairing LEC-autophagy causes an overall accumulation of naïve T cells in SLOs. Furthermore, although to a lesser extent, we found a trend of increased B cell (CD3^-^ CD19^+^) and NK cell abundance (CD3^-^ NK1.1^+^) in the SLOs of ATG5^LEC-KO^ mice compared with their WT counterparts (Supplementary Fig. 1 D-F). Despite the overall increase of T cell, B cell, and NK cell numbers, their frequencies were not altered by the deletion of ATG5 in LECs (Supplementary Fig. 1G). Also, the size and weight of inguinal LN and spleen were not significantly increased (Supplementary Fig. 1H,I).

**Figure 1.**
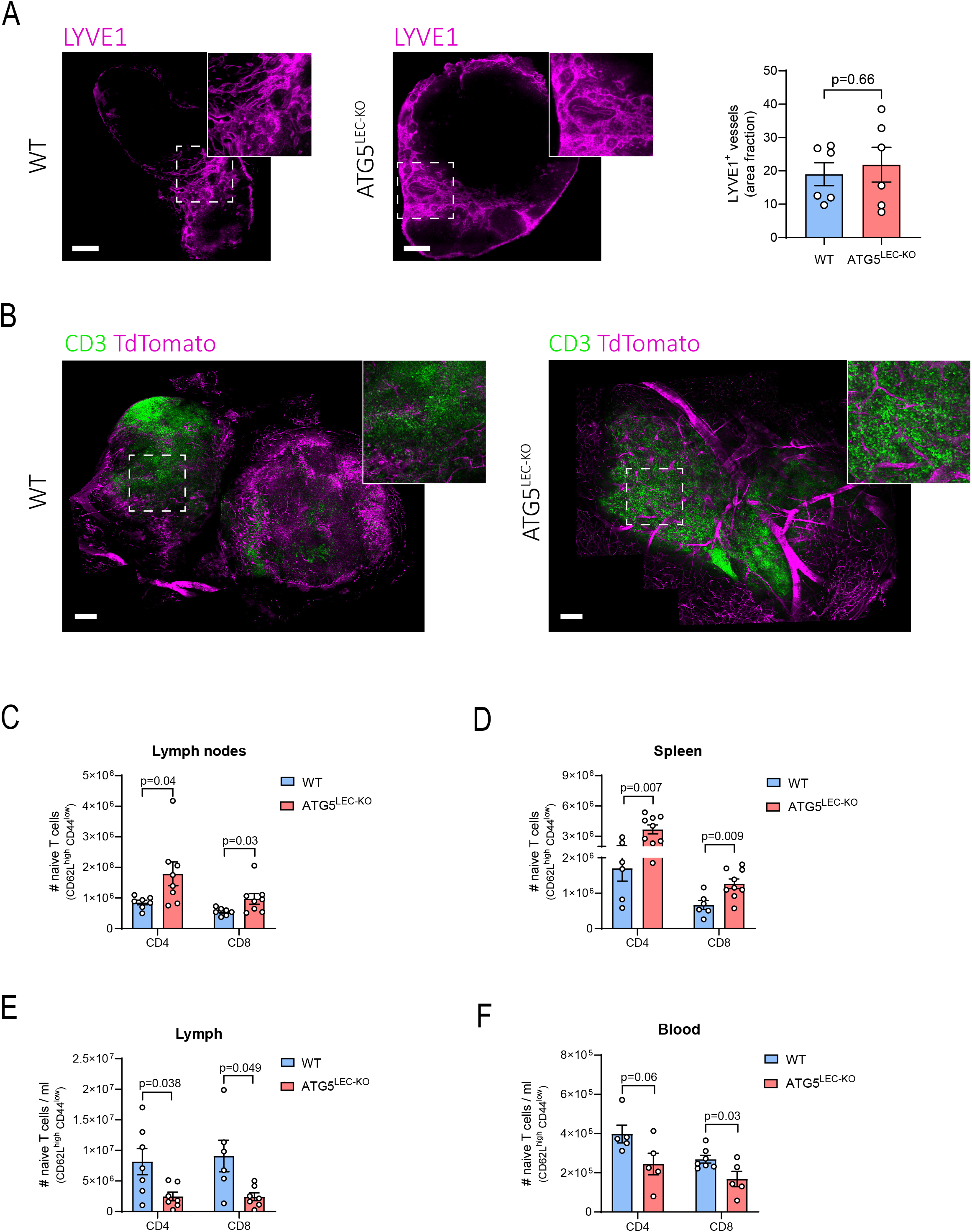
Genetic deletion of ATG5 in LECs impairs the recirculation of lymphocytes in secondary lymphoid organs at steady-state. **A)** Representative immunofluorescent images, corresponding zooms of region of interest (ROI), and quantification of LYVE1 area fraction (magenta) of whole inguinal lymph nodes from wild-type (WT) and ATG5^LEC-KO^ mice at steady-state. Mean ± SEM, n=6 inguinal lymph nodes from independent mice analyzed using unpaired t-test. Scale bars 200 µm. **B)** Representative 3D immunofluorescent images and corresponding crop of cleared whole inguinal lymph nodes from wild-type (WT) and ATG5^LEC-KO^ mice harboring widespread tdTomato fluorescence expression (magenta) in cells/tissues at steady- state. Lymph nodes were stained for CD3 (green). Scale bars 200 µm. **C)** Total number of naïve (CD62L^high^ CD44^low^) CD4 and CD8 T cells in pooled lymph nodes from wild-type (WT) and ATG5^LEC-KO^ mice at steady-state. Mean ± SEM, each data point represents one mouse, n=7 analyzed using Mann-Whitney test. **D)** Number of naïve (CD62L^high^ CD44^low^) CD4 and CD8 T cells in spleens from wild-type (WT) and ATG5^LEC-KO^ mice at steady-state. Mean ± SEM, each data point represents one mouse, n=6 analyzed using unpaired t-test. **E)** Total number of naïve (CD44^low^ CD62L^high^) CD4 and CD8 T cells per ml of lymph fluid from wild-type (WT) and ATG5^LEC-KO^ mice at steady-state. Mean ± SEM, each data point represents one mouse, n≥6 analyzed using unpaired t-test with Welch’s correction. **F)** Numbers of naïve (CD62L^high^ CD44^low^) CD4 and CD8 T cells per ml of blood from wild-type (WT) and ATG5^LEC-KO^ mice at steady-state. Mean ± SEM, each data point represents one mouse, n≥5 analyzed using unpaired t-test.

A previous report attributed fluctuations in T cell number in SLOs to changes in T cell survival (28). To study this possibility, we analyzed cell death (PI staining) and proliferation (Ki67) of naïve CD4 and CD8 T cells isolated from LNs of WT and ATG5^LEC-KO^ mice. We found no major differences in the fraction of dying or proliferative T cell populations between these conditions and although significant, the decreased CD4 T cell proliferation was of minor magnitude (Supplementary Fig. 2A,B). A possible mechanism explaining the persuasive phenotype observed in ATG5^LEC-KO^ could be a defect in the trafficking of lymphocytes between LNs and lymph, which would also manifest in lymphopenia. To this end, we collected lymph fluid directly from the cisterna chyli, while blood samples were collected from the submandibular vein. Compared with WT mice, ATG5^LEC-KO^ mice showed a comparable trend towards fewer naïve T cells (Fig. 1E,F), B cells, and NK cells (Supplementary Fig. 2C,D) in both the lymphatic and blood circulatory systems.

Together, these data indicate that loss of LEC-autophagy leads to the sequestration of major lymphocyte populations in SLOs.

### Autophagy in LECs controls T cell egress from lymph nodes under steady-state conditions

Next, we sought to gain further insights into the mechanisms of disturbed T cell redistribution caused by the deletion of ATG5 in LECs. First, we analyzed whether the increase in lymphocyte number in LNs of ATG5^LEC-KO^ micewas facilitated by structural changes or alterations in the number of HEVs, the main entry point for blood-borne lymphocytes into the LN, via adherence of the T cell integrins LFA-1 and VLA-4 to endothelial ligands (29). Immunostaining for HEVs with MECA79, an antibody that recognizes the HEV-specific peripheral node addressin (PNAd) (29,30), did not reveal apparent changes in the typical cuboidal structure of HEVs, nor in their number or density when comparing LNs from WT or ATG5^LEC-KO^ mice (Supplementary Fig. 3A-C). We then examined whether the accumulation of lymphocytes in the LN of the LEC-ATG5 deficient mice was caused by an enhanced entry, which could point to functional changes in the HEVs, or by a defect of lymphocytes egress, which would increase their permanence in the LN parenchyma. To evaluate these conjectures, we divided WT and ATG5^LEC-KO^ mice into two cohorts and quantified naïve CD4 T and CD8 T cells from their LNs, immediately (time zero, first cohort) or after 8h following blockade of T cell entry by the intravenous injection of both LFA-1 and VLA-4 neutralizing antibodies (time 8 hours, second cohort) (31) (Fig. 2A). We then calculated the percentage of CD4 and CD8 T cells remaining in the LNs of these mice. After 8 hours of T cell entry blockade, the LN of WT mice contained around 35% of the initial population of CD4 and CD8 T cells, whereas at the same time point, nearly double the fraction (e.g. 60-70 %) of T cells was retained in the LN of ATG5^LEC-KO^ mice (Fig. 2B). Together these findings pointed to a defect in T cell egress from the LN of LEC-autophagy deficient mice.

**Figure 2.**
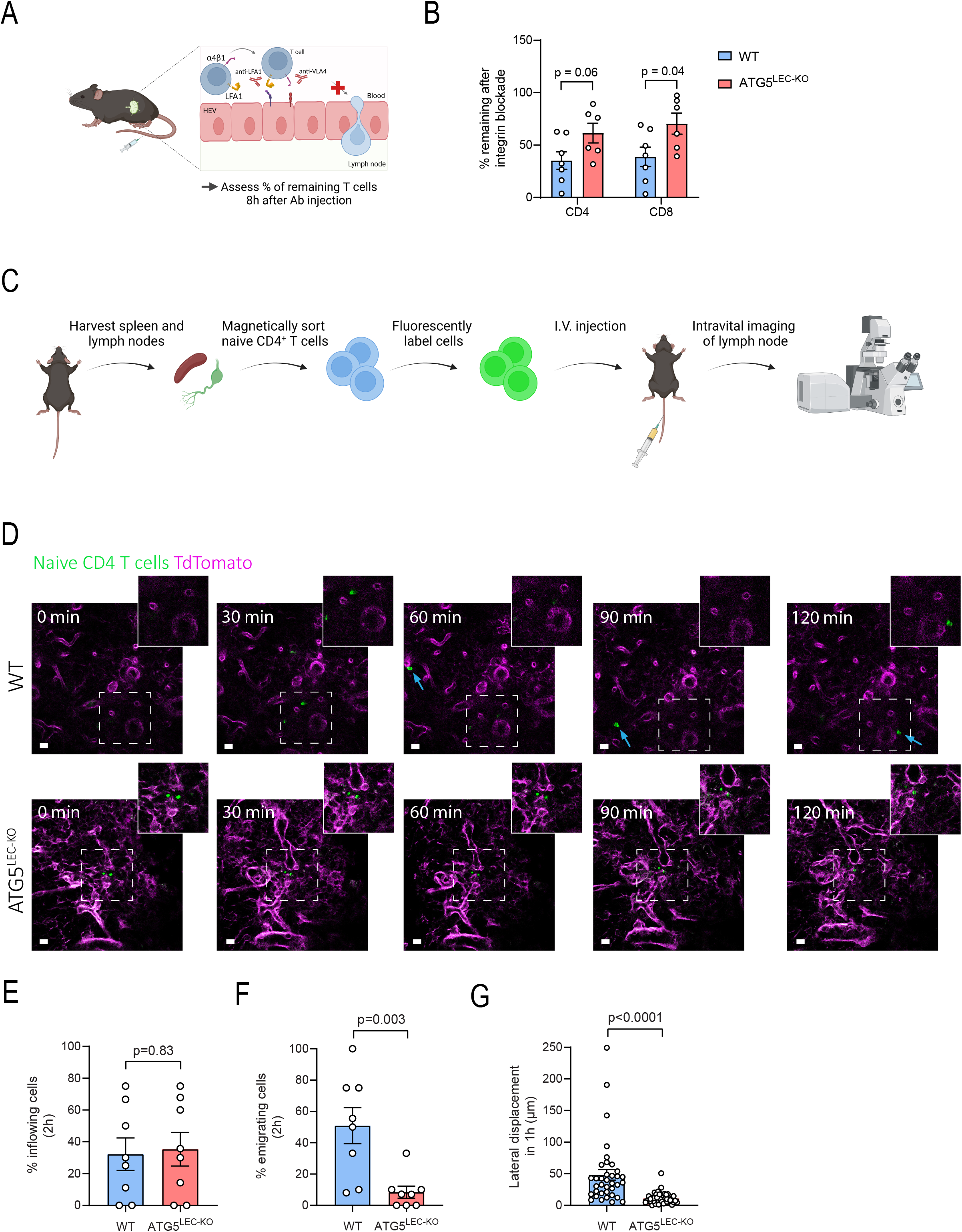
Loss of ATG5 in LECs causes a defect in T cell egress from lymph nodes. **A)** Experimental design of the experiments in B. Mice were injected intravenously with 100 μg monoclonal antibody to integrin α4 and 100 μg monoclonal antibody to integrin αL and sacrificed eight hours later. **B)** Percentage of remaining CD4 or CD8 T cells in pooled lymph nodes 8 h after integrin blockade with anti-LFA-1 and anti-VLA-4 antibodies (as compared to the average number in LNs at t=0h) of wild-type (WT) and ATG5^LEC-KO^ mice. Each point represents one mouse at t = 8h relative to the average at t = 0h. Mean ± SEM, n≥6 analyzed using unpaired t-test. **C)** Experimental design of the experiments in D. Spleens and LNs were harvested and naïve CD4 T cells were magnetically sorted and fluorescently labeled with a Biotracker 655 dye. Two million cells were injected intravenously and intravital imaging started two hours later. **D)** Representative images and corresponding crop of different time points of intravital imaging of inguinal lymph nodes of wild-type (WT) and ATG5^LEC-KO^ mice. Timepoint “0 min” corresponds to 2 hours after intravenous injection of labeled CD4 naïve T cells (green). Scale bars 10 µm. **E)** Percentage of inflowing naïve CD4 T cells in the inguinal lymph nodes over a period of two hours in wild-type (WT) and ATG5^LEC-KO^ mice. Mean ± SEM, each data point represents one area (ROI) analyzed from a total of 8 mice from 4 independent experiments using unpaired t-test. **F)** Percentage of emigrating naïve CD4 T cells that move away in the inguinal lymph nodes over a period of two hours in wild-type (WT) and ATG5^LEC-KO^ mice. Mean ± SEM, each data point represents one area analyzed from a total of 8 mice from 4 independent experiments using Mann-Whitney test. **G)** Lateral displacement of naïve CD4 T cells in the inguinal lymph nodes over a period of one hour in wild-type (WT) and ATG5^LEC-KO^ mice. Images of different time points (1h interval) representing the same Z-stack were merged and displacement was measured by drawing a line between the same cell throughout the different time points. Mean ± SEM, each data point represents one cell analyzed from a total of 8 mice from 4 independent experiments using Mann-Whitney test.

To further monitor the intranodal dynamics of this process in the live setting, we utilized intravital time-lapse imaging of the inguinal LN using a skin flap in WT and ATG5^LEC-KO^ R26-mTmG mice. As reported in previous studies (32) we used magnetically sorted naïve CD4 T cells that were fluorescently labeled with the Biotracker 655 Red cytoplasmic membrane dye and injected intravenously to allow their visualization within the LN (Fig. 2C). To image T cell entry into the LN, we started the intravital imaging two hours after T cell injection and followed T cell dynamics over the next 2-3 hours (Fig. 2D, Supplementary Fig. 3D). To visualize the entry dynamics of adoptively transferred T cells, we focused on areas enriched in HEVs, likely representing the paracortex, as confirmed by intravenous injection of a conjugated MECA79 antibody (Supplementary Fig. 3E). In WT mice, T cells were initially found in near HEVs and moved away during the 2 hour imaging period, following a known random walking behavior (Fig. 2D, Video 3) (33). In contrast, while the percentage of inflowing naïve T cells was similar in ATG5^LEC-KO^ mice, the percentage of T cells that emigrated from the entry site was drastically reduced (Fig. 2E,F, Video 4). Upon arrival in the LN, T cells of ATG5^LEC-KO^ mice displayed a static behavior with limited lateral displacement over 1 hour, compared to the exhibiting motility of T cells injected in WT mice (Fig. 2G). These data suggest a disturbed balance between the retention and exit signals to which T cells are exposed within the LN parenchyma, resulting in their impaired egress.

Together, these results indicate that LEC-autophagy promotes the egress of naïve T cells from the LNs, thereby maintaining constitutive lymphocyte recirculation between SLOs and the blood.

### Naïve CD4/CD8 T cells in SLOs of ATG5^LEC-KO^ mice are exposed to higher levels of S1P

The egress of naïve T cells from the LN is primarily regulated by the S1P gradient, which is mainly controlled by LECs in the LN (9). To define whether ATG5^LEC-KO^ mice displayed unbalanced extracellular S1P levels compared with WT mice in the LN, we analyzed the surface abundance of the S1P receptor 1 (S1PR1) on naïve T cells by FACS, which is considered the gold-standard methodology, validated in different studies to measure ‘signaling-available’ S1P. Naïve T cells sensing low S1P levels have high S1PR1 levels on their surface, while naïve T cells sensing high S1P levels have lower surface levels of the S1PR1 receptor as it gets internalized and partially degraded upon S1P binding (Fig. 3A) (9,34).

**Figure 3.**
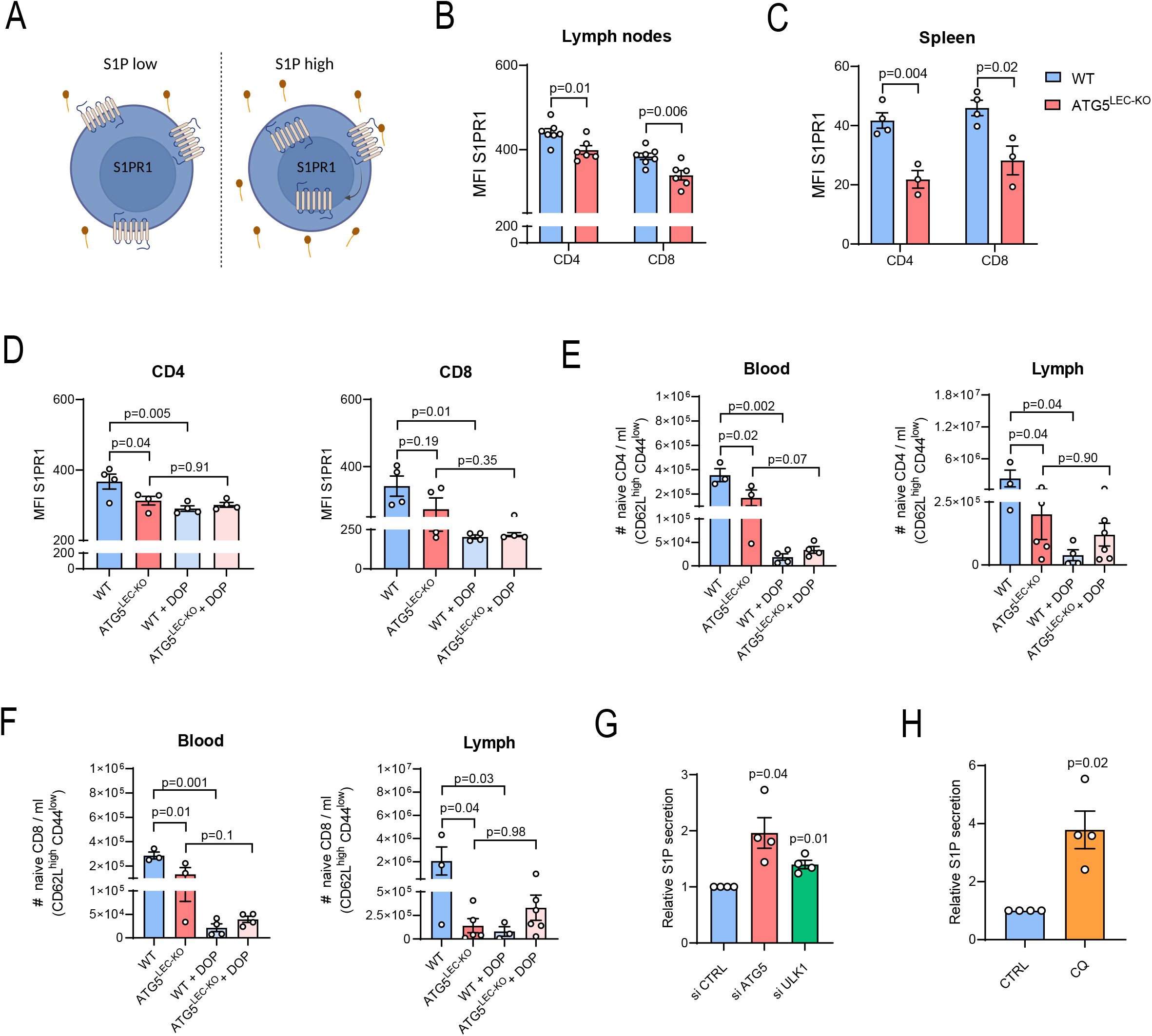
Naïve CD4/CD8 T cells in the SLOs of ATG5^LEC-KO^ mice are exposed to higher levels of S1P. **A)** Illustration of the ‘signaling-available’ S1P measurement *in vivo*. Upon low encounter of S1P, naïve T cells express S1PR1 on the surface. When exposed to high levels of S1P, the S1PR1 receptor gets internalized after binding, leading to its reduced surface levels. **B)** Median fluorescence intensity (MFI) of S1PR1 on naïve CD4 and CD8 T cells in pooled lymph nodes from wild-type (WT) and ATG5^LEC-KO^ mice at steady-state. Mean ± SEM, each data point represents one mouse, n≥6 analyzed using unpaired t-test. **C)** Median fluorescence intensity (MFI) of S1PR1 on naïve CD4 and CD8 T cells in spleens from wild-type (WT) and ATG5^LEC-KO^ mice at steady-state. Mean ± SEM, each data point represents one mouse, n≥3 analyzed using unpaired t-test. **D)** Median fluorescence intensity (MFI) of S1PR1 on naïve CD4 and CD8 T cells in pooled lymph nodes from wild-type (WT) and ATG5^LEC-KO^ mice treated with DOP. Mean ± SEM, each data point represents one mouse, n=4 analyzed by one-way ANOVA corrected for multiple comparisons using Dunn’s test. **E)** Numbers of naïve (CD62L^high^ CD44^low^) CD4 T cells per ml of lymph fluid or blood from wild-type (WT) and ATG5^LEC-KO^ mice treated with DOP. Mean ± SEM, each data point represents one mouse, n≥3 analyzed by one-way ANOVA corrected for multiple comparisons using Sidak’s test. **F)** Numbers of naïve (CD62L^high^ CD44^low^) CD8 T cells per ml of lymph fluid or blood from wild-type (WT) and ATG5^LEC-KO^ mice treated with DOP. Mean ± SEM, each data point represents one mouse, n≥3 analyzed by one-way ANOVA corrected for multiple comparisons using Sidak’s test. **G)** Relative S1P secretion in siCTRL, siATG5, and siULK1 Human dermal lymphatic endothelial cells (HDLECs) in 10% lipid-stripped serum medium after 8h. Mean ± SEM, n=4 biological replicates analyzed using one sample t-test. **H)** Relative S1P secretion in CTRL or CQ-treated HDLECs in 10% lipid stripped serum medium after 8h. Mean ± SEM, n=4 biological replicates analyzed using one sample t-test.

When comparing the S1PR1 levels on both naïve CD4 and CD8 T cells from pooled LNs and spleens of WT or ATG5^LEC-KO^ mice by FACS, we observed that naïve T cells of ATG5^LEC-KO^ mice had reduced S1PR1 surface levels, thus suggesting their exposure to higher S1P levels within these SLOs (Fig. 3B,C). The increase in intranodal S1P levels could flatten the S1P gradient in homeostatic LNs, thereby abolishing the directional exit signal for naïve T cells. To confirm the relevance of the S1P signaling axis, we treated WT or ATG5^LEC-KO^ mice with 4-deoxypyridoxine (DOP), a known inhibitor of the S1P-degrading enzyme S1P lyase, which has been reported to increase intranodal S1P levels, thereby preventing T cell egress (12)(28). Congruently, DOP treatment reduced S1PR1 surface levels on naïve T cells in LNs of WT mice, but did not induce a further reduction in ATG5^LEC-KO^ mice (Fig. 3D, Supplementary Fig. 4A). In addition, DOP treatment reduced the number of circulating naïve T cells in WT mice to a similar extent as observed in ATG5^LEC-KO^ mice, while it failed to further decrease their number in ATG5^LEC-KO^ mice, suggesting a similar mode of action (Fig. 3E,F).

T cell egress from the thymus occurs via blood vessels in an S1P-S1PR1-dependent manner (11). Neither the total number of single-positive thymocytes, cell frequencies, or thymocyte maturation were changed when comparing WT with ATG5^LEC-KO^ mice (Supplementary Fig. 4B-E), thus indicating that the effects of loss of LEC-autophagy on the S1P-S1PR1 axis were lymph-specific.

To gain mechanistic insights, we assayed whether autophagy blockade in LECs increased the secretion of S1P *in vitro,* using primary human dermal LECs (HDLECs). Silencing ATG5 or ULK1, a component of the early autophagosome formation complex (35), or treatment with chloroquine (CQ), an inhibitor of the autophagosome-lysosome fusion (36), resulted in enhanced secretion of S1P by HDLECs (Fig. 3G,H). To further understand the mechanism driving this increase in S1P secretion, we characterized expression levels of the proteins involved in S1P production and secretion. S1P kinase 1 (SPHK1), catalyzes the conversion of sphingosine to S1P and the S1P transporter SPNS2 is known to be essential in S1P secretion in lymph (37). While neither SPHK1 nor SPNS2 protein or RNA levels were altered in autophagy-depleted HDLECs, we observed an accumulation of serine palmitoyl transferase (SPT), the rate-limiting enzyme in de novo synthesis of sphingolipids (Supplementary Fig. 4F-I). SPT is an ER-localizing enzyme (Supplementary Fig. 4J) previously shown to be an autophagic cargo (38). In line with this, while loss of autophagy did not alter significantly *SPT* expression (Supplementary Fig. 4F,G), blockade of lysosomal degradation by CQ increased SPT colocalization with the autophagosomal marker LC3, indicating that SPT undergoes constitutive autophagic recycling (Supplementary Fig. 4K), likely through a process known as ER-phagy (39).

Together, these results suggest that loss of autophagy in LECs increases intranodal S1P levels, thereby impairing the main exit signal for naïve T cells.

### LEC-mediated trafficking and positioning of lymphocytes in tumor-draining LNs requires ATG5

Given the pervasive T cell trafficking defect caused by loss of ATG5 in LECs under steady-state conditions, we next asked whether this effect was phenocopied in the TdLNs of tumor-bearing mice. To this end, we subcutaneously inoculated B16F10 melanoma cells in WT or ATG5^LEC-KO^ mice and when the tumor reached a size of maximum 1500 mm^3^ (day 17, endpoint), we isolated inguinal TdLNs, identified by intratumoral injection of Evans Blue (Supplementary Fig. 5A). While tumor growth in WT and ATG5^LEC-KO^ mice was similar (Fig. 4A,B), the TdLNs of ATG5^LEC-KO^ mice showed a significant increase in T cell cellularity (Supplementary Fig. 5B), a minor decrease in lymphatic vessel density (Supplementary Fig. 5C) and harbored an elevated number of naïve CD4 and CD8 T cells when compared with TdLNs from WT mice (Fig. 4C). We next evaluated whether T cells in the TdLNs of melanoma-bearing ATG5^LEC-KO^ mice sensed higher levels of S1P. Congruent with the phenotype observed under steady-state conditions, S1P levels were heightened in the TdLNs of ATG5^LEC-KO^ mice (Fig. 4D). Similar results were obtained using the HCMel12-mCherry melanomas, carrying *Hgf* and *Cdk4r244c* mutations (Supplementary Fig. 5D-G), suggesting that the phenotype driven by the loss of *Atg5* in LECs is independent of the melanoma genetic background. Importantly, TdLNs from mice harboring Atg5 deletion in blood ECs (ATG5^BEC-KO^), obtained by crossing *Atg5^fl/fl^* mice with *Pdgf-cre^ERT2^*mice (40), did not show an increase in the cellularity of the CD4 and CD8 naïve T cell population nor a change in S1PR1 surface levels (Supplementary Fig. 5H,I). Hence, the effects of Atg5 on the dynamics of T cells in the TdLNs are LEC-specific.

**Figure 4.**
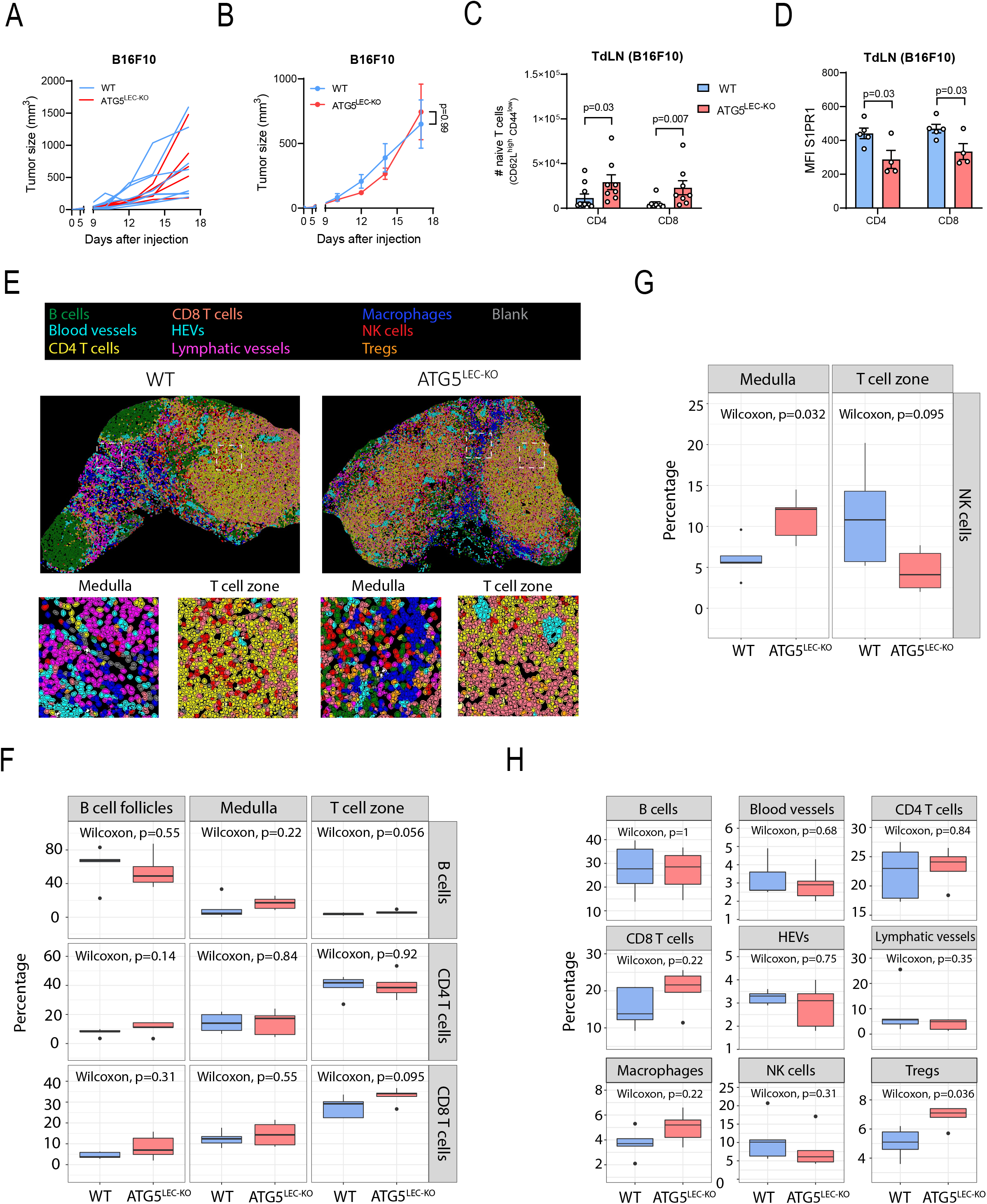
Loss of ATG5 in LECs alters T cell trafficking in the TdLNs of tumor-bearing mice. **A)** Tumor growth curve of individual wild-type (WT) and ATG5^LEC-KO^ mice injected with B16F10 tumors. N=7 and5 respectively. **B)** Tumor growth curve of wild-type (WT) and ATG5^LEC-KO^ mice injected with B16F10 cells. Mean ± SEM, n=5-7 mice per group analyzed using two-way ANOVA corrected for multiple comparisons using Sidak’s test. **C)** Number of naïve (CD62L^high^ CD44^low^) CD4 and CD8 T cells in TdLNs from wild-type (WT) and ATG5^LEC-KO^ mice bearing B16F10 tumors. Mean ± SEM, each data point represents one mouse, n≥6 analyzed using Mann-Whitney test. **D)** Median fluorescence intensity (MFI) of S1PR1 on naïve CD4 and CD8 T cells in the TdLNs from wild-type (WT) and ATG5^LEC-KO^ mice bearing B16F10 tumors. Mean ± SEM, each data point represents one mouse, n≥4 analyzed using unpaired t-test. **E)** Representative digital reconstruction of the tissue based on multiplex staining and cell clustering illustrating EC subtypes and immune cells in the TdLNs with crops of the medulla and T cell zone of wild-type (WT) and ATG5^LEC-KO^ mice bearing B16F10 tumors. **F)** Distribution of B cells, CD4 and CD8 T cells in B cell follicles, medulla, and T cell zones of the TdLNs of wild-type (WT) and ATG5^LEC-KO^ mice bearing B16F10 tumors. Mean ± SEM, n=5 analyzed using two-sided Wilcoxon test. **G)** Distribution of NK cells in the medulla and T cell zones of the TdLNs of wild-type (WT) and ATG5^LEC-KO^ mice bearing B16F10 tumors. Mean ± SEM, n=5 analyzed using two-sided Wilcoxon test. **H)** Percentage of cells from indicated populations on total cells in the TdLNs of wild-type (WT) and ATG5^LEC-KO^ mice bearing B16F10 tumors. Mean ± SEM, n=5 analyzed using two-sided Wilcoxon test.

Within the highly compartmentalized LNs, lymphocyte distribution is organized in a tightly controlled manner. Although the exact S1P concentration throughout the LN parenchyma is unknown, alterations in the S1P gradient may result in the intranodal relocation of lymphocytes (41). To gain insights into the anatomical positioning of immune cells within the TdLN parenchyma of WT and ATG5^LEC-KO^ mice, we utilized multiple iterative labeling by antibody neo-deposition (MILAN) multiplex immunohistochemistry (42,43). This allowed us to annotate different immune cell clusters, including CD4 and CD8 T cells, Tregs, B cells, NK cells, and macrophages, alongside major vessel types within TdLNs, including HEVs, lymphatic vessels, and blood vessels (Fig. 4E, Supplementary Fig. 5J,K). We then focused on the positioning of lymphocytes within the medulla, B cell follicles and the T cell zone of TdLNs of tumor-bearing WT and ATG5^LEC-KO^ mice (Supplementary Fig. 5L). Both B cells and T cells did not show overall displacements, although TdLNs of ATG5^LEC-KO^ showed a trend towards an increased CD8 T cellularity within the T cell zones (Fig. 4F). Interestingly, deletion of LEC-ATG5 induced a trend toward an increase in the B cells and CD8 T cells in the T zone, and a marked relocation of NK cells from the T cell zone towards the medullary regions of TdLN parenchyma (Fig. 4E,G), in line with a recent study describing NK cell delocalization upon intranodal S1P gradient defects (41). Compared with WT mice, the fractions of the main population of lymphocytes, macrophages, and endothelial cells on total cells (Fig. 4H) did not show major variations in TdLNs of the ATG5^LEC-KO^ mice, in line with our immune profiling and immunofluorescent analysis. The minor fraction of Tregs was elevated in TdLNs of the ATG5^LEC-KO^ mice (Fig. 4H), likely because of an exacerbated S1P- S1PR1 axis, which has been shown to promote the differentiation of Foxp3 Tregs (44,45).

Thus, LEC-autophagy orchestrates the spatiotemporal presence of lymphocytes in the TdLN of melanoma-bearing mice.

### Single-cell transcriptomics identifies distinct ATG5-regulated signatures in LN LEC niches

The changes in T cell trafficking and NK cell zonation observed in the TdLNs of the ATG5^LEC-KO^ mice suggest that deletion of ATG5 may elicit a transcriptional and functional reprogramming of LECs. To map LEC heterogeneity and identify potential processes regulating lymphocyte migratory function and LEC-T cell interactions, at single-cell resolution, we performed scRNAseq analysis of inguinal TdLNs from B16F10 tumor-bearing WT and ATG5^LEC-KO^ mice. To increase the coverage of different LEC subtypes, we pooled a total of 13 and 9 TdLNs from melanoma-bearing WT and ATG5^LEC-KO^ mice, respectively. We then FACS-sorted viable CD45^-^ Ter119^-^ CD31^+^ endothelial cells (ECs) and CD45^+^ Ter119^-^ CD31^-^ immune cells, yielding a total of 12500 cells, and processed them for scRNAseq analysis using the BD Rhapsody platform (Fig. 5A). After quality and cell filtering, we retained a total of 3118 ECs and 5732 CD45+ immune cells in our analysis. Subsequently, we performed unsupervised clustering for ECs and immune cells, according to their gene-expression profiles, based on the expression of known markers and used Uniform Manifold Approximation and Projection (UMAP) for visualization. Using curated canonical markers, we identified distinct subsets of adaptive and innate immune cells and the two main EC subtypes, which included HEVs and LECs (Fig. 5B, Supplementary Fig. 6A). As expected, the autophagy gene signature in LECs was downregulated upon genetic deletion of *Atg5* in LECs (selected from the Reactome), confirming the shutdown of autophagy pathways at the transcriptional level (Supplementary Fig. 6B).

**Figure 5.**
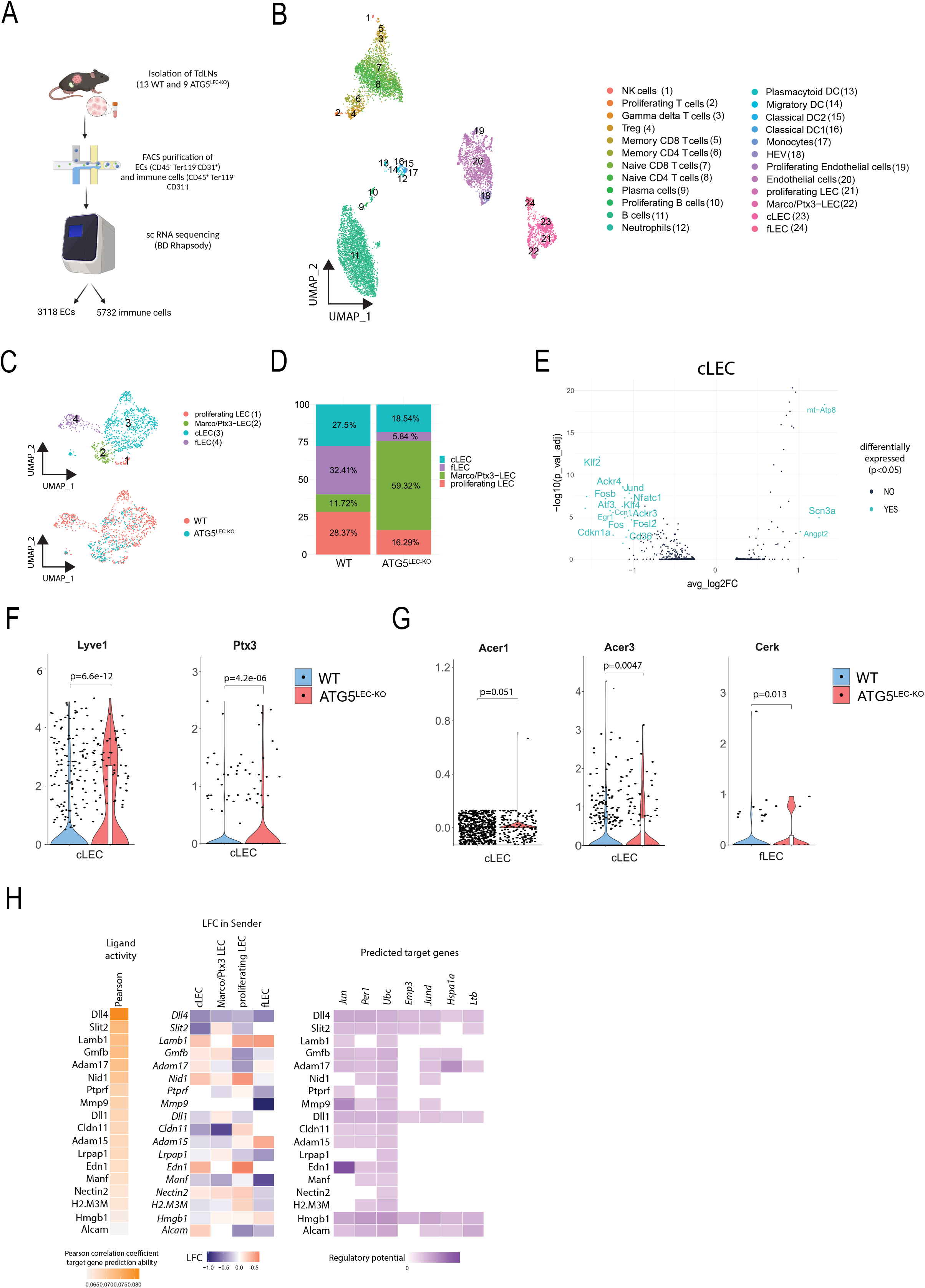
scRNAseq of TdLN reveals niche-specific changes in LEC LN subsets. **A)** Illustration of the experimental set-up. **B)** UMAP embedding plot of 8850 LN-resident cells grouped into 24 clusters. **C)** UMAP embedding plot of LN-resident LECs visualized into 4 clusters and per condition. **D)** Barplot of LEC subtype frequencies of total LECs in the TdLNs of B16F10 tumor-bearing WT and ATG5^LEC-KO^ mice. **E)** Volcano plot showing differentially expressed genes between LN cLECs from WT and ATG5^LEC-KO^ mice. P value < 0.05 & log2 fold change > 1 or < −1, analyzed using two-sided Wilcoxon test. **F)** Violin plot showing expression of respective “gene” across cLECs in TdLNs of B16F10 tumor-bearing WT and ATG5^LEC-KO^ mice. Median-quantile-min/max + population distribution is shown in violin and boxplot analyzed using two-sided Wilcoxon test. **G)** Violin plot showing expression of respective “gene” across different subtypes of LECs in TdLNs of B16F10 tumor-bearing WT and ATG5^LEC-KO^ mice. Median-quantile-min/max + population distribution is shown in violin and boxplot analyzed using two-sided Wilcoxon test. **H)** Outcome of NicheNet’s ligand activity prediction: results are shown for the top 18 LEC-ligands. (from left to right) Pearson correlation indicates the ability of each ligand to predict the target genes, and better predictive ligands are thus ranked higher. Expression of ligands in different LEC subtypes. Log fold change expression of ligands comparing WT and ATG5^LEC-KO^ LEC subtypes. NicheNet’s ligand–target matrix denotes the regulatory potential between LEC ligands and target genes.

Subclustering of LECs, identified by consistent expression of LEC markers *Nr2f2/Prox1*/*Pdpn,* resulted in distinct subpopulations, in line with previous reports (15,17), including (i) ceiling LEC (cLEC), characterized by the expression of *Ackr4/Gdf10/Jag1* with low *Lyve1* levels, (ii) floor LEC (fLEC) expressing *Bmp2/Csf1/Smad4/Vcam1*, (iii) Marco/Ptx3-LEC expressing *Lyve1/Flt4/Nrp2/Ptx3/Cd81* and (iv) a small cluster of proliferating LECs, marked by high levels of *Mki67/Stmn1/Cenpe/Cdk1,* sharing markers with both cLEC and Marco/Ptx3 LECs (Supplementary Fig. 6A).

All subtypes were present in both WT and ATG5^LEC-KO^ samples. Of note, looking at the compositional alterations in LECs, at the subcluster level between WT and ATG5^LEC-KO^ mice samples, we observed changes in niche-associated LEC abundancies, indicating a reduction in the proportions of the cLEC and fLEC subtypes lining the subcapsular sinus (SCS) lumen, facing the outside or the parenchyma respectively, and an enrichment in the *Lyve1*+ Marco/Ptx3-LEC cluster in ATG5^LEC-KO^ TdLNs (Fig. 5C,D). We then performed differential gene expression (DEG) analysis, using a strict threshold (cut off p < 0.05 & log2FC > 1 or < −1) to characterize the distinctive features of the LEC subtypes and the functional processes influenced by the loss of ATG5.

We identified cLECs as the cluster whose transcriptional profile was most prominently affected by the loss of ATG5, based on the abundance of DEG with significant effect sizes (Fig. 5E, Supplementary Fig. 6C). While few genes were upregulated (e.g. *Angpt2* implicated in VEGFR3-mediated lymphangiogenesis (46)), deletion of ATG5 caused a dominant downregulation of several genes involved in cLEC proliferation (*Egr1, Cdkn1a, Klf2)*, VEGF-C/VEGFR3-driven PROX1 expression (*Nfatc1, Klf4*, *Ccn1)* and immediate early transcription factors downstream of VEGFR3/VEGFR2 (*Jund, FosB, Fos, Fosl2, Atf3*), suggesting a pro-lymphangiogenic action of ATG5 in cLECs (Fig. 5E). Congruently, single-cell regulatory network inference and clustering (SCENIC) (47) analysis showed downregulation of *Jun*, *Egr1* and *Fos*-related regulons in ATG5-deficient cLECs, further indicating that loss of ATG5 remodels gene expression signatures, particularly in the cLEC subtype (Supplementary Fig. 6D).

Interestingly, conforming to recent scRNAseq data in both mice and humans (15,17), specific cLEC markers, including the atypical chemokine receptor *Ackr4/CCRL1*, which regulates the physiological gradient of CCL19 and CCL21 from SCS into the LN parenchyma (48), and the fatty acid transporter *CD36* involved in oxLDL uptake (49), were also downregulated by ATG5 deletion, suggesting a loss of cLEC specification. In line, we observed that cLECs from ATG5^LEC-KO^ mice tended to increase the expression of *Lyve1* and *Ptx3,* two distinctive markers of Marco-Ptx3-LECs (Fig. 5F), further implicating the tendency to acquire a more ‘medullary-like’ phenotype, with increased capability to control lymphocytes egress (15–17). Congruently, immunofluorescent staining for LYVE1 and ACKR4 in TdLNs of WT and ATG5^LEC-KO^ mice revealed the presence of a distinguishable ACKR4+/LYVE1- vessel layer of cLECs decorating the SCS in WT mice, which became less distinct in the ATG5^LEC-KO^ mice, suggesting the gain of a more ACKR4-/LYVE1+ cLEC phenotype following the loss of ATG5 (Supplementary Fig. 6E).

Our results portray that impaired lymphocyte egress from TdLNs of the ATG5^LEC-KO^ mice is caused by enhanced secretion of the lipid S1P. Thus, we focused on the expression of genes involved in lipid metabolism and in particular, sphingolipid signaling, known to be implicated in the production of S1P in the identified LN LEC subtypes. Specifically, we observed that the ER-Golgi-associated ceramidases (*Acer1, Acer3)* were enriched in cLECs and ceramide kinase (*Cerk)* in fLECs (Fig. 5G), further suggesting a transcriptional alteration in key components of the pathway leading to increased production of S1P by these LEC subtypes.

Increased permanence of lymphocytes in the TdLN of ATG5^LEC-KO^ mice (50) could also manifest in the alteration of their transcriptional profile. Using established markers of memory, naïve cells, and regulatory T cells (Tregs) for CD4 T cells, we found that CD4 and CD8 T cell subclusters did not exhibit significant compositional alterations between TdLNs from WT and LEC-specific Atg5 KO mice (Supplementary Fig. 6F,G). However, cell-cell communication analysis with NicheNet (51) revealed potentially perturbed LEC-T cell bidirectional interaction pathways between WT and ATG5^LEC-KO^ mice. NicheNet predicted the following ligands, expressed by LECs, as the most likely regulators of DE genes between T cells from WT and LEC-ATG5 deficient mice; the Notch1/2 ligands, *Dll4, Nid1, Adam17, Dll1,* suggesting that LECs act as nonhematopoietic sources of Notch ligands in the TdLN, with potential effects on T cell differentiation and fate (52,53), and *Slit2/Lamb1* involved in T cell chemotaxis/directionality (Fig. 5H) (54,55). Interestingly, the expression of several top predicted LEC ligands, was affected following the loss of ATG5 in a LEC-subtype specific manner. In particular, while following the deletion of Atg5, *Dll4* and *Lamb1* expression was respectively up- or downregulated in all LEC LN subtypes, *Slit2* expression was reduced in cLECs, while being upregulated in Marco/Ptx3 LECs, further suggesting perturbations in molecular signals regulating T cell recruitment and directionality (Fig. 5H).

Thus, LEC-ATG5 is required to maintain the cLEC transcriptional phenotype and niche-specific signals predicted to regulate LEC-T cell molecular interactions.

### Loss of LEC-ATG5 abrogates the anti-tumor immunity effect of immunotherapy

Recent studies highlight that the recruitment of peripheral T cells to the tumor plays a critical role in the efficacy of immunotherapies (19,20). We then hypothesized that ICB-mediated therapy would be less effective in melanoma-bearing ATG5^LEC-KO^ mice compared to WT mice. Congruent with this hypothesis, in B16F10 tumor-bearing WT mice, anti-PD-1 therapy reduced tumor burden and weight and led to an increase in tumor-infiltrating lymphocytes (TILs), in particular CD8 T cells and NK cells (Fig. 6A-F, Supplementary Fig. 7A). In contrast, anti-PD-1 therapy failed to increase TILs and control tumor burden in melanoma-bearing ATG5^LEC-KO^ mice (Fig. 6A-F, Supplementary Fig. 7A), but it elevated the number of CD4, CD8 T cells and NK cells in the TdLN of WT mice to a similar extent to what observed in TdLN of ATG5^LEC-KO^ mice (Supplementary Fig. 7B). Notwithstanding, the composition of TILs (Supplementary Fig. 7C-E) and the fraction of CD8 T cells expressing the known activation markers, Granzyme B, KLRG1, and IFNγ and the proliferation marker Ki67 (Fig. 6G,H, Supplementary Fig. 7F-H) were unaltered in melanomas grown in WT and ATG5^LEC-KO^ mice. Together, these results suggest that ICB-driven antitumor responses fail in melanoma-bearing ATG5^LEC-KO^ mice, because of a defect in the migration of ICB-responsive (tumor-reactive) lymphocytes to the tumor, which is manifested in a decrease in their absolute number. To corroborate this hypothesis, we co-treated melanoma-bearing mice with anti-PD-1 and the S1P lyase inhibitor DOP, which elevated S1P levels in the LN (Fig. 3D). Blockade of lymphocyte egress by DOP abrogated tumor response mediated by anti-PD-1 therapy in melanoma-bearing WT mice, but failed to exert any additional effects in the ATG5^LEC-KO^ mice (Fig. 6I-K). Hence, the blockade of lymphocyte exit through the dysregulation of the S1P gradient in the LNs likely explains how the loss of LEC-autophagy neutralizes the antitumor immunity effects of ICBs.

**Figure 6.**
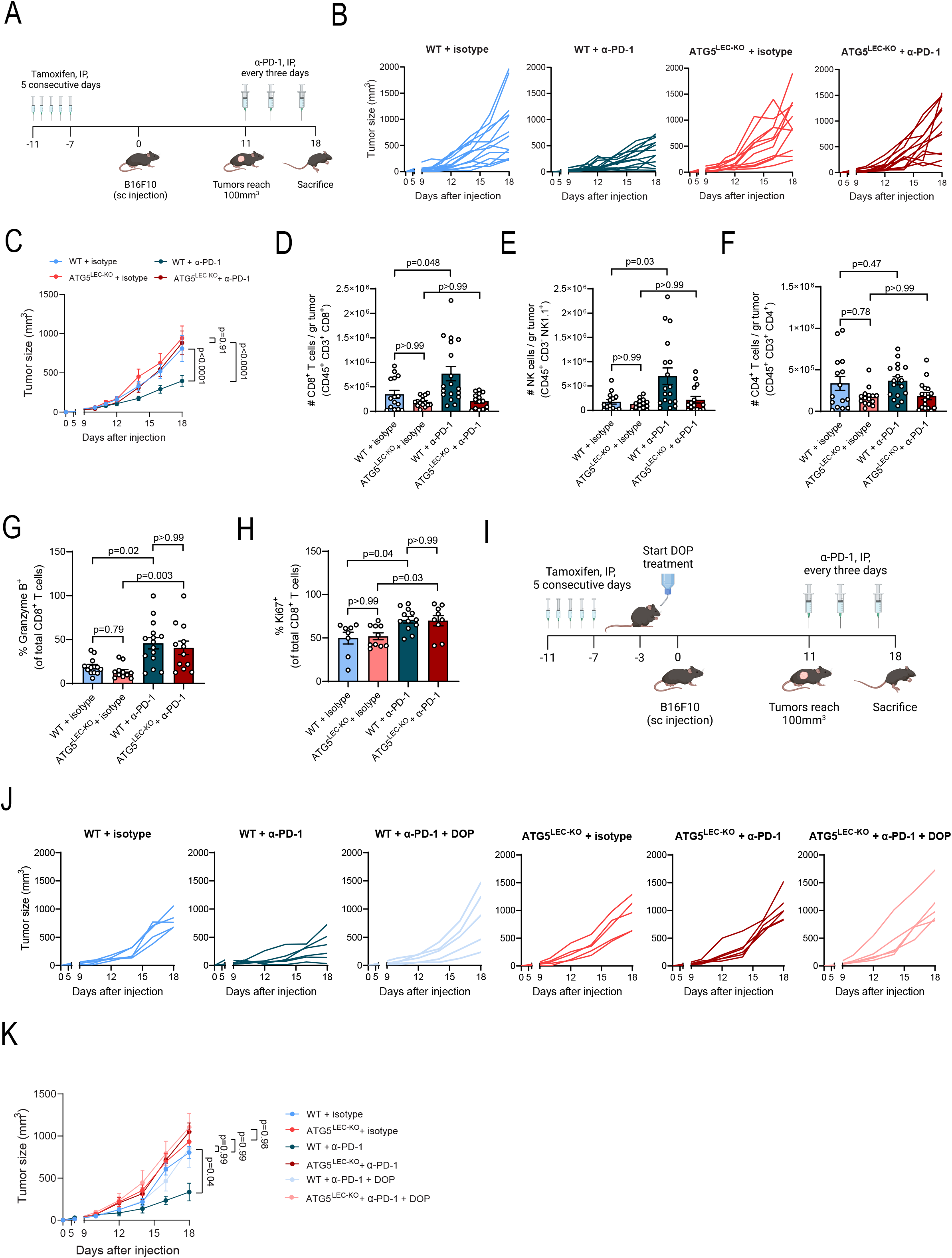
Loss of LEC-ATG5 blocks the anti-tumor effect of ICB. **A)** Experimental design of B-H. Mice were subcutaneously injected with 150.000 B16F10 cells. Once the tumors reached 100mm3, 250µg of α-PD-1 mAb or 250 µg of isotype was injected intraperitoneally twice a week. **B)** Tumor growth curve of individual wild-type (WT) and ATG5^LEC-KO^ mice injected with B16F10 cells treated with 250µg isotype antibody or 250µg α-PD-1 antibody twice per week starting the moment tumors reached 100mm^3^, n=13, 13, 11, 12 respectively. **C)** Tumor growth curve of wild-type (WT) and ATG5^LEC-KO^ mice injected with B16F10 cells treated with 250µg isotype antibody or 250µg α-PD-1 antibody twice per week starting the moment tumors reached 100mm^3^, n=11-13 mice per group analyzed using two-way ANOVA corrected for multiple comparisons using Sidak’s test. **D)** Number of intratumoral CD8 T cells (CD45^+^ CD3^+^ CD8^+^) of wild-type (WT) and ATG5^LEC-KO^ mice from A. Mean ± SEM, each data point represents one mouse, n≥16 mice per group analyzed using Kruskal-Wallis test corrected for multiple comparisons using Dunn’s test. **E)** Number of intratumoral NK cells (CD45^+^ CD3^-^ NK1.1^+^) of wild-type (WT) and ATG5^LEC-KO^ mice from A. Mean ± SEM, each data point represents one mouse, n≥14 mice per group analyzed using Kruskal-Wallis test corrected for multiple comparisons using Dunn’s test. **F)** Number of intratumoral CD4 and CD8 T cells in HCMel12-mCherry tumors. Mean ± SEM, each data point represents one mouse, n≥5 analyzed using unpaired t-test (CD8) or Mann-Whitney test (CD4). **G)** Frequency of intratumoral Granzyme B^+^ CD8 T cells on total CD8 T cells of wild-type (WT) and ATG5^LEC-KO^ mice from A. Mean ± SEM, each data point represents one mouse, n≥11 mice per group analyzed using Welch ANOVA test corrected for multiple comparisons using Dunnett’s T3 test. **H)** Frequency of intratumoral Ki67^+^ CD8 T cells of total CD8 T cells of wild-type (WT) and ATG5^LEC-KO^ mice from A. Mean ± SEM, each data point represents one mouse, n≥8 mice per group analyzed using Kruskal-Wallis test corrected for multiple comparisons using Dunn’s test. **I)** Experimental design of J,K. **J)** Tumor growth curve of individual wild-type (WT) and ATG5^LEC-KO^ mice injected with B16F10cells treated with 250µg isotype antibody or 250µg α-PD-1 antibody twice per week starting the moment tumors reached 100mm^3^ and treated with DOP. n=5, 6, 5, 5, 5, 5 respectively. **K)** Tumor growth curve of wild-type (WT) and ATG5^LEC-KO^ mice injected with B16F10 cells treated with 250µg isotype antibody or 250µg α-PD-1 antibody twice per week starting the moment tumors reached 100mm^3^ and treated with DOP. Mean ± SEM, n≥5 mice per group analyzed using two-way ANOVA corrected for multiple comparisons using Tukey’s test.

In a previous study, immunotherapy responses in the poor lymphangiogenic melanoma model B16F10 were improved by potentiating peritumoral lymphangiogenesis, through the overexpression of VEGF-C in B16F10 cells (56). In B16F10VEGF-C melanomas, an increased influx of CCR7-expressing naïve T cells undergoing local activation after immunotherapy overruled the heightened immunosuppressive tumor microenvironment caused by the infiltration of myeloid cells and Tregs in the untreated tumors (56). We then asked whether the phenotype observed in B16F10-bearing ATG5^LEC-KO^ mice would be altered in this more lymphangiogenic model. B16F10VEGF-C melanomas in WT mice, as expected, displayed a higher density of Lyve1+/CD31+ positive lymphatic vessels compared with the poor lymphangiogenic B16F10 tumors (Supplementary Fig. 8A) and increased tumor burden (Supplementary Fig. 8B,C). In mice harboring B16F10VEGF-C melanomas, both tumor burden and peritumoral lymphangiogenesis (Supplementary Fig.8A-C) were reduced by the loss of Atg5 in LECs. Concurrently, the increased infiltration of various inflammatory and immunosuppressive cells observed in previous studies (56), was dampened in B16F10VEGF-C tumor-bearing ATG5^LEC-KO^ mice compared with their WT counterparts (Supplementary Fig. 8D). In contrast, no significant changes were observed in the non-lymphangiogenic B16F10 melanomas (Supplementary Fig. 8E). Notwithstanding, the TdLNs from the B16F10VEGF-C melanomas of ATG5^LEC-KO^ mice displayed significant retention of naïve CD4 and CD8 T cells, hallmarked by the surface downregulation of the S1PR1 (Supplementary Fig. 8F,G) and a higher number of NK cells (Supplementary Fig. 8H).

Remarkably, while anti-PD-1 therapy induced effective tumor regression in B16F10VEGF-C-bearing WT mice, it failed to elicit antitumor responses and abrogated the tumor burden control observed in ATG5^LEC-KO^ mice (Supplementary Fig. 8I,J). Notably, also, in the ICB-responsive MC38 colon carcinoma model, loss of LEC-ATG5 blunted tumor regression after anti-PD-1 therapy (Fig. 7A,B). Although we cannot exclude the contribution of additional tumor-intrinsic mechanisms, together, these results suggest that the loss of ATG5 in LECs abrogates antitumor immunity induced by ICB largely by impairing the trafficking of ICB-responsive T lymphocytes. Furthermore, loss of ATG5 in LECs curbed tumor regression after anti-CTLA4 immunotherapy (Fig. 7C,D), which is thought to act primarily at the site of T cell priming, thus the LNs, by enhancing CD28 costimulation (57). This finding further indicates that loss of ICB responsiveness in tumor-bearing ATG5^LEC-KO^ mice is independent of the ICB treatment modality.

**Figure 7.**
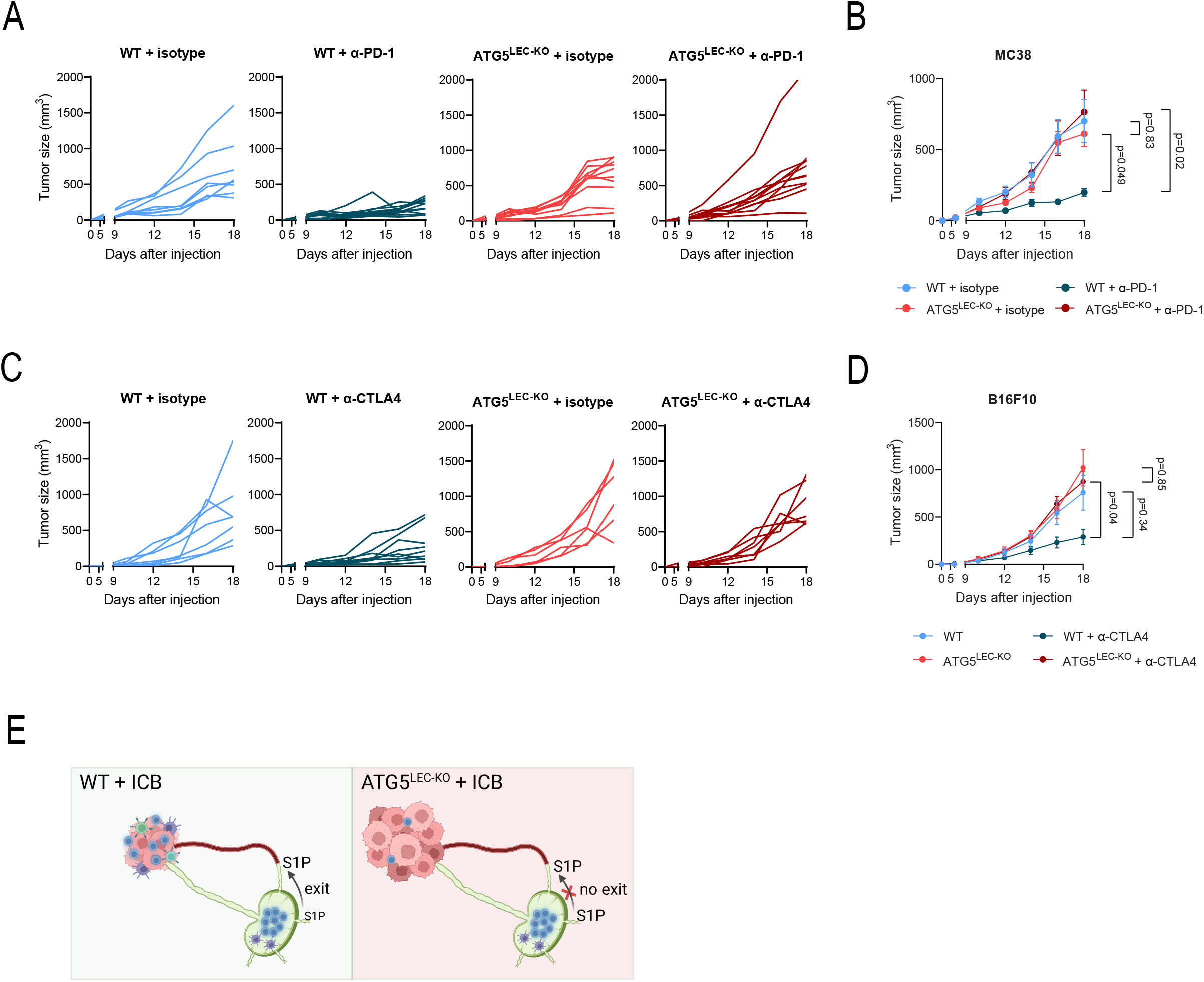
ICB resistance in ATG5^LEC-KO^ mice is independent of the tumor model or ICB modality. **A)** Tumor growth curve of individual wild-type (WT) and ATG5^LEC-KO^ mice injected with MC38 cells treated with 250µg isotype antibody or 250µg α-PD-1 antibody twice per week starting the moment tumors reached 100mm^3^, n=8,10,12,11 respectively. **B)** Tumor growth curve of wild-type (WT) and ATG5^LEC-KO^ mice injected with MC38 cells treated with 250µg isotype antibody or 250µg α-PD-1 antibody twice per week starting the moment tumors reached 100mm^3^. Mean ± SEM, n=8-12 mice per group analyzed using two-way ANOVA corrected for multiple comparisons using Sidak’s test. **C)** Tumor growth curve of individual wild-type (WT) and ATG5^LEC-KO^ mice injected with B16F10 tumors treated with 100µg isotype antibody or 100µg α-CTLA4 antibody twice per week starting the moment tumors reached 100mm^3^. n=7, 9, 6, 7 respectively. **D)** Tumor growth curve of wild-type (WT) and ATG5^LEC-KO^ mice injected with B16F10 tumors treated with 100µg isotype antibody or 100µg α-CTLA4 antibody twice per week starting the moment tumors reached 100mm^3^. Mean ± SEM, n=6-9 mice per group analyzed using two-way ANOVA corrected for multiple comparisons using Sidak’s test. **E)** Illustration of the working model of immune checkpoint blockade resistance in ATG5^LEC-KO^ mice.

In aggregate, our results show that the antitumor immunity effects of the most prominent ICB molecules require functional autophagy in LN LECs, which orchestrates the trafficking of T cells from the periphery to the tumor (Fig. 7E).

## DISCUSSION

A growing body of evidence indicates that SLOs and peripheral T cell dynamics are critical determinants of T cell responses to immunotherapy (18–21). Thus, gaining comprehensive knowledge of the mechanisms controlling global immune dynamics is mandatory to improve our understanding of how immunotherapy works and how we can ameliorate its efficacy. Our study identifies a fundamental autophagy-regulated program in LECs that controls the recirculation dynamics of T cells from the LNs to the tumor, which is a prerequisite for the efficacy of immunotherapies.

Previous studies have proposed that host autophagy represses antitumor immunity mainly through its innate ability to remove major sources of endogenous and extracellular inflammatory signals (58). However, the functional outcome of autophagy pathways is highly cell-type and stress-specific and thus inevitably complex. Here, by combining dynamic intravital imaging, immune profiling and high-resolution multiplex imaging of major lymphocytes populations, we found that autophagy in LECs of the LNs regulates lymphocyte compartmentalization and migration within the LN parenchyma, by regulating the availability of the major egress signal S1P. Only recently, single-cell profiling of LECs has started to unravel the molecular heterogeneity and niche-specific specialization of lymphatic vessels of the lymph nodes. However, the molecular mechanisms that control the distinct niche-specific molecular phenotypes of LN LECs are unknown. Through our unbiased single-cell sequencing data analysis of TdLNs, we portrayed that LECs lining the ceiling of the SCS are particularly reliant on autophagy to maintain their transcriptional phenotype. Within the SCS, the site where lymphatic vessels enter the LN capsule, cLECs play a role in guiding leukocyte attraction to the LN parenchyma through the formation of a CCL21 gradient, a function requiring the expression of *Ackr4* (48). Therefore, the downregulation of the cLEC-specific markers, *Ackr4* and *Cd36,* and the enrichment in the more medullary/paracortical markers (e.g. *Lyve1, Ptx3*) of the PTX3-LECs involved in lymphocytes’ exit route (17), following the deletion of ATG5, is particularly intriguing and suggests that regulation of lymphocytes trafficking within LN is distinctively altered by the loss of autophagy. Interestingly, following optogenetic stimulation of sensory neurons innervating the SCS of murine LNs, cLECs showed selective downregulation in the expression of genes involved in the turnover of S1P, suggesting a role for SCS innervation in lymphocytes egress (59). Whether the function of autophagy in cLECs is controlled by peripheral neurons, innervating the LN is an interesting perspective requiring future studies.

Notwithstanding, the exact reason for the selective autophagy dependence of cLECs remains elusive. Because of their specific LDL scavenging role, cLECs could be more prone to alterations in lipid homeostasis, a process that is fine-tuned by autophagy (60,61). We recently reported that in a model of injury-driven corneal lymphangiogenesis, LEC-specific deletion of ATG5 impairs lipophagy (the specific degradation of lipid droplets by autophagy) and the mitochondria fatty acid oxidation-derived production of acetyl-CoA, which fosters PROX1-driven transcription in LECs (25). ATG5 deletion in LECs of the LNs, and in particular in the cLEC subtype, however, does not result in the downregulation of direct PROX1 targets, such as VEGFR3. Conceivably, the functional outcome of autophagy pathways in LECs reflects specific molecular adaptations to inflammation-inducing factors, the composition of the lymph, and the highly specialized microenvironment of the LN, characterized by high levels of fatty acids and a distinct profile of cytokines and chemokines. Consistent with this, recent scRNAseq studies unraveled the unique transcriptomic signatures and functional and molecular heterogeneity of LECs from different anatomical localizations (62,63). The molecular adaptations and transcriptional heterogeneity of LECs could also explain why in a previous study, ATG5 deletion in LN LECs aggravated collagen-induced arthritis, by promoting the egress of pathogenic Th17 cells from the LN to the inflamed joint (27).

A full understanding of the relative contribution of local and systemic antitumor immunity and their dynamic crosstalk during immunotherapy is still lacking. Our study provides compelling evidence showing that mice lacking LEC-specific autophagy do not respond to the two major clinically used immunotherapies, anti-PD-1 and anti-CTLA4. This is largely because following ICB, primed T cells fail to traffic from the periphery to the tumor parenchyma. In line, TILs in ATG5^LEC-KO^ mice do not display gross alterations in their activation status after immunotherapy, highlighting that it is the reduction in their absolute number that ablates T cell response and consequent ICB-mediated tumor regression. Our data however do not exclude that other signals regulated by LEC-autophagy within the TME may further contribute to the reduced number of TILs observed in the tumor-bearing ATG5^LEC-KO^ mice. Notwithstanding, the results of our study show that the effects of loss of LEC-autophagy on T cell dynamics are largely uncoupled from the inflammatory status of the TME, since in all tumor models tested, ICB failed to induce tumor regression, further indicating their systemic immunity nature. Consistent with this notion, recent studies both in humans and mice have shown that TdLNs act as reservoirs of ICB-responsive T cells and that tumor-reactive T cells may be activated peripherally and then recruited to the TME, where they acquire their canonical effector function (21,64–66). Selective targeting of the PD-L1/PD-1 axis only in TdLN has been shown to generate effective antitumor immune responses (67). Finally, other reports have shown that FTY720 treatment may abolish the effect of ICB, although the effect of this S1P analog is dependent on the onset of the treatment (20,56,68,69), further suggesting that early T cell recruitment towards the tumor site is essential for ICB efficacy. Our experiments did not examine clonal T cell dynamics, which require TCR sequencing studies from both TdLN and tumor-derived T lymphocytes. Future work should address the initial priming steps within the TdLN and identify which ‘precursor’ T cells are mainly affected in terms of egress in mice with LEC-specific loss of autophagy.

Our preclinical data demonstrate that functional autophagy signaling in LECs of the LNs is required for the therapeutic efficacy of ICB, providing further molecular insights into the relevance of LN immune-vascular interaction and its consequence for immunotherapy. While the direct therapeutic implications of our findings require further exploration, the results of this study advocate for the use of targeted autophagy modulators or interventions as a mechanism to amplify peripheral T cell recruitment to the tumor. Here, we show that genetic loss of *Atg5* in blood ECs (BECs) does not phenocopy the effects on T cell dynamics observed in the TdLN of ATG5^LEC-KO^ mice. However, we recently found that genetic loss of autophagy in blood ECs endorses NF-kB and STING signaling in the tumor vasculature of melanoma-bearing mice, which correlated with anti-PD-1 therapy responses, by supporting local T cell infiltration and effector function (70). Hence, future therapeutic interventions that aim to harness autophagy in combination with immunotherapies need to consider the functional impact of this catabolic pathway on the immunomodulatory functions of ECs from different vascular beds, which our studies have revealed. Vasculature-homing tools, potentially via tissue-specific nanobodies, may allow autophagy modulation within the preferred vascular bed, without exerting additional unwanted effects on others.

## LEGENDS

**Supplementary Figure 1.**

**A)** Representative immunofluorescent images and corresponding crop of LYVE1^+^ lymphatic vessels (magenta) in lymph nodes from wild-type (WT) and ATG5^LEC-KO^ mice stained for ATG5 (green) at steady-state. Scale bars 50 µm. **B)** Representative confocal immunofluorescent images of LYVE1^+^ lymphatic vessels (magenta) in lymph nodes from wild-type (WT) and ATG5^LEC-KO^ mice stained for LC3B (green) at steady-state. Arrows indicate LC3B puncta. Scale bars 10 µm. **C)** Representative immunofluorescent images and quantification of CD3+ area fraction(green) of whole lymph nodes from wild-type (WT) and ATG5^LEC-KO^ mice at steady-state. Nuclei are stained with DAPI (blue). Mean ± SEM, n≥6 lymph nodes from independent mice analyzed using unpaired t-test. Scale bars 200 µm. **D)** Gating strategies for naïve T cells, B cells, and NK cells. **E)** Number of B cells (CD3^-^ CD19^+^) and NK cells (CD3^-^ NK1.1^+^) in pooled lymph nodes from wild-type (WT) and ATG5^LEC-KO^ mice at steady-state. Mean ± SEM, each data point represents one mouse, n≥8 analyzed using unpaired t-test (B cells) with Welch’s correction (NK cells). **F)** Number of B cells (CD3^-^ CD19^+^) and NK cells (CD3^-^ NK1.1^+^) in spleens from wild-type (WT) and ATG5^LEC-KO^ mice at steady-state. Mean ± SEM, each data point represents one mouse, n≥8 analyzed using unpaired t-test (B cells) with Welch’s correction (NK cells). **G)** Frequency of CD4 and CD8 T cells, B cells, and NK cells of total CD45^+^ live singlets in pooled lymph nodes from wild-type (WT) and ATG5^LEC-KO^ mice at steady-state. Mean ± SEM, each data point represents one mouse, n≥6 analyzed using unpaired t-test (B and NK cells) with Welch’s correction (CD4 and CD8). **H)** Representative images and weight of inguinal lymph nodes from wild-type (WT) and ATG5^LEC-KO^ mice at steady-state. Mean ± SEM, n=4 lymph nodes from independent mice analyzed using unpaired t-test. Scale bar represents 5 mm. **I)** Representative images and weight of spleens from wild-type (WT) and ATG5^LEC-KO^ mice at steady-state. Mean ± SEM, n=7 spleens from independent mice analyzed using unpaired t-test. Scale bar represents 10 mm.

**Supplementary Figure 2.**

**A)** Frequency of propidium iodide (PI) positive naïve CD4 and CD8 T cells in lymph nodes from wild-type (WT) and ATG5^LEC-KO^ mice at steady-state. Mean ± SEM, each data point represents one mouse, n=6 analyzed using Mann-Whitney test. **B)** Frequency of proliferating (Ki67^+^) naïve CD4 and CD8 T cells in lymph nodes from wild-type (WT) and ATG5^LEC-KO^ mice at steady-state. Mean ± SEM, each data point represents one mouse, n≥6 analyzed using unpaired t-test (CD4) with Welch’s correction (CD8). **C)** Number of B cells (CD3^-^ CD19^+^) and NK cells (CD3^-^ NK1.1^+^) per ml of lymph fluid from wild-type (WT) and ATG5^LEC-KO^ mice at steady-state. Mean ± SEM, each data point represents one mouse, n≥7 analyzed using unpaired t-test (B cells) with Welch’s correction (NK cells). **D)** Number of B cells (CD3^-^ CD19^+^) and NK cells (CD3^-^ NK1.1^+^) per ml of blood from wild-type (WT) and ATG5^LEC-KO^ mice at steady-state. Mean ± SEM, each data point represents one mouse, n≥6 analyzed using Mann-Whitney test.

**Supplementary Figure 3.**

**A)** Representative immunofluorescent images and corresponding crop of MECA79^+^ lymphatic vessels (magenta) in the lymph nodes from wild-type (WT) and ATG5^LEC-KO^ mice at steady-state. Nuclei are stained with DAPI (blue). Scale bars 100 µm. **B)** Number of MECA79^+^ vessels per mm^2^ in the lymph nodes from wild-type (WT) and ATG5^LEC-KO^ mice at steady-state. Mean ± SEM, each data point represents one mouse, n≥5 analyzed using unpaired t-test. **C)** Area fraction of MECA79^+^ vessels in the lymph nodes from wild-type (WT) and ATG5^LEC-KO^ mice at steady-state. Mean ± SEM, each data point represents one mouse, n≥4 analyzed using unpaired t-test with Welch’s correction. **D)** Representative images and corresponding crop of different time points of intravital imaging in the inguinal lymph nodes of wild-type (WT) and ATG5^LEC-KO^ mice. Timepoint “0 min” corresponds to 2 hours after intravenous injection of labeled CD4 naïve T cells (green). Scale bars 10 µm. **E)** Representative intravital images of inguinal lymph nodes of wild-type (WT) R26-mTmG mice 2h after injection of fluorescently labeled CD4 naïve T cells and MECA79-conjugated antibodies. Scale bar represents 10 µm.

**Supplementary Figure 4.**

**A)** Representative plot of surface S1PR1 on naïve CD4 T cells in pooled lymph nodes from wild-type (WT) and ATG5^LEC-KO^ mice treated with DOP. **B)** Number of single positive CD4 and CD8 T cells in the thymus from wild-type (WT) and ATG5^LEC-KO^ mice at steady-state. Mean ± SEM, each data point represents one mouse, n≥7 analyzed using unpaired t-test (CD4) with Welch’s correction (CD8). **C)** Frequency of single positive and double positive (DP) CD4 and CD8 T cells of total live singlets (SP) in the thymus from wild-type (WT) and ATG5^LEC-KO^ mice at steady-state. Mean ± SEM, each data point represents one mouse, n=9 analyzed using unpaired t-test (DP) or Mann-Whitney test (CD4 SP and CD8 SP). **D)** Number of mature single positive CD4 and CD8 T cells (CD69^-^ CD62L^high^) in the thymus from wild-type (WT) and ATG5^LEC-KO^ mice at steady-state. Mean ± SEM, each data point represents one mouse, n≥3 analyzed using unpaired t-test. **E)** Frequency of mature single positive CD4 and CD8 T cells (CD69^-^ CD62L^high^) in the thymus from wild-type (WT) and ATG5^LEC-KO^ mice at steady-state. Mean ± SEM, each data point represents one mouse, n≥4 analyzed using unpaired t-test with Welch’s correction. **F)** RT-qPCR analysis of siCTRL, siATG5, and siULK1 HDLECs. mRNA expression of SPTLC1, SPTLC2, SPNS2 and SPHK1 (relative to HPRT). Mean ± SEM, n=3 biological replicates, p=0.50, p=0.26, p=0.71, p=0.41 (siATG5) and p=0.09, p=0.20, p=0.74, p=0.05 (siULK1) analyzed using one sample t-test. **G)** RT-qPCR analysis of CTRL or CQ-treated HDLECs. mRNA expression of SPTLC1, SPTLC2, SPNS2 and SPHK1 (relative to HPRT). Mean ± SEM, n=3 biological replicates, p=0.28, p=0.67, p=0.18, p=0.77 analyzed using one sample t-test. **H)** Representative blots and densitometric quantification of the indicated proteins in siCTRL, siATG5, and siULK1 HDLECs. Mean ± SEM, n=3 biological replicates analyzed using one sample t-test. **I)** Representative blots and densitometric quantification of the indicated proteins in CTRL or CQ-treated HDLECs. Mean ± SEM, n=4 biological replicates analyzed using one sample t-test. **J)** Representative confocal immunofluorescent images of LEC stained for PDI as ER-marker (green) and SPT (magenta). Nuclei are stained with DAPI. Scale bars 10µm. **K)** Representative confocal immunofluorescent images and colocalization analysis (Manders M1 coefficient) of CTRL or CQ-treated LEC after mCherry-LC3 transfection (green) stained for SPT (magenta). Nuclei are stained with DAPI. n= 30 cells from 3 biological replicates, analyzed by Mann-Whitney test. Scale bars 10 µm.

**Supplementary Figure 5.**

**A)** Representative image of a tumor-draining lymph node (TdLN) and non-TdLN identified after intratumoral Evans blue injection. Scale bar represents 1 mm. **B)** Representative immunofluorescent images and quantification of CD3+ area fraction (green) in the TdLNs from wild-type (WT) and ATG5^LEC-KO^ mice bearing HCMel12-mCherry tumors. Nuclei are stained with DAPI. Scale bars 100 µm. Mean ± SEM, n≥7 lymph nodes from independent mice analyzed using unpaired t-test. **C)** Representative immunofluorescent images, corresponding crop, and quantification of LYVE1 area fraction (magenta) in the TdLNs from wild-type (WT) and ATG5^LEC-KO^ mice bearing HCMel12-mCherry tumors. Nuclei are stained with DAPI. Scale bars 100 µm. Mean ± SEM, n=8 lymph nodes from independent mice analyzed using Mann-Whitney test. **D)** Tumor growth curve of individual wild-type (WT) and ATG5^LEC-KO^ mice injected with HCMel12-mCherry cells. n=12. **E)** Tumor growth curve of wild-type (WT) and ATG5^LEC-KO^ mice injected with HCMel12-mCherry cells. Mean ± SEM, n=12 mice per group analyzed using two-way ANOVA corrected for multiple comparisons using Sidak’s test. **F)** Number of naïve (CD44^low^ CD62L^high^) CD4 and CD8 T cells in tumor-draining lymph nodes from wild-type (WT) and ATG5^LEC-KO^ mice bearing HCMel12-mCherry tumors. Mean ± SEM, each data point represents one mouse, n≥6 analyzed using unpaired t-test with Welch’s correction. **G)** Median fluorescence intensity (MFI) of S1PR1 on naïve CD4 and CD8 T cells in tdLNs from wild-type (WT) and ATG5^LEC-KO^ mice bearing HCMel12-mCherry tumors. Mean ± SEM, each data point represents one mouse, n≥4 analyzed using unpaired t-test with Welch’s correction (CD4 HCMel12-mCherry) or Mann-Whitney test (CD8 HCMel12-mCherry). **H)** Number of naïve (CD44^low^ CD62L^high^) CD4 and CD8 T cells in tumor-draining lymph nodes from wild-type (WT) and ATG5^BEC-KO^ mice bearing HCMel12mCherry tumors. Mean ± SEM, each data point represents one mouse, n≥4 analyzed using unpaired t-test. **I)** Median fluorescence intensity (MFI) of S1PR1 on naïve CD4 and CD8 T cells in the TdLNs from wild-type (WT) and ATG5^BEC-KO^ mice bearing HCMel12mCherry tumors. Mean ± SEM, each data point represents one mouse, n≥4 analyzed using unpaired t-test with Welch’s correction. **J)** Heatmap showing markers used for the annotation of cell populations using MILAN. **K)** Representative digital reconstruction of the tissue based on multiplex staining and cell clustering illustrating EC subtypes and immune cells in the TdLNs of wild-type (WT) and ATG5^LEC-KO^ mice bearing B16F10 tumors. **L)** Representative digital reconstruction of the tissue based on multiplex staining with the demarcation of indicated zones within the TdLNs.

**Supplementary Figure 6.**

**A)** A dotplot showing marker genes across cell clusters from the UMAP embedding plot in Figure 6A. The data point size indicates the percent of expressed cells, and the color represents the average expression level. **B)** Heatmap of Reactome signatures for autophagy-related pathways in all LEC from TdLNs from wild-type (WT) and ATG5^LEC-KO^ mice. **C)** Volcano plot showing differentially expressed genes between LN fLECs and Marco/Ptx3-LECs from wild-type (WT) and ATG5^LEC-KO^ mice. P value < 0.05 & log2 fold change > 1 or < −1, analyzed using two-sided Wilcoxon test. **D)** Heatmap of selected inferred transcription-factor gene-regulatory networks (SCENIC). **E)** Representative confocal immunofluorescent images and corresponding zooms of region of interest (ROI) of LYVE1 fluorescence intensity in ACKR4+ lymphatic vessels (green) in TdLNs from wild-type (WT) and ATG5^LEC-KO^ mice stained for LYVE1 (magenta) at steady-state. Scale bars 10µm.**F)** UMAP embedding plot of LN-resident T and NK cells grouped into 8 clusters per condition. **G)** Barplots of CD4 and CD8 T cell subtype frequencies of total CD4 and CD8 T cells in TdLNs of B16F10 tumor-bearing WT and ATG5^LEC-KO^ mice.

**Supplementary Figure 7.**

**A)** B16F10 tumor weight at day 18 after tumor cell injection. Mean ± SEM, each data point represents one mouse, n≥16-18 per group analyzed using one-way ANOVA corrected for multiple comparisons using Holm-Sidak’s test. **B)** Number of CD8 T cells (CD45^+^ CD3^+^ CD8^+^), NK cells (CD45^+^ CD3^-^ NK1.1^+^), and CD4 T cells (CD45^+^ CD3^+^ CD4^+^) in the TdLNs of wild-type (WT) and ATG5^LEC-KO^ mice injected with B16F10 cells treated with 250µg isotype antibody or 250µg α-PD-1 antibody twice per week starting the moment tumors reached 100mm^3^. Mean ± SEM, each data point represents one mouse, n≥14 mice per group analyzed using Kruskal-Wallis test corrected for multiple comparisons using Dunn’s test. **C)** Frequency of intratumoral CD8 T cells (CD45^+^ CD3^+^ CD8^+^) of total CD45^+^ cells of wild-type (WT) and ATG5^LEC-KO^ mice injected with B16F10 cells treated with 250µg isotype antibody or 250µg α-PD-1 antibody twice per week starting the moment tumors reached 100mm^3^. Mean ± SEM, each data point represents one mouse, n≥16 mice per group analyzed using one-way ANOVA test corrected for multiple comparisons using Holm-Sidak’s test. **D)** Frequency of intratumoral CD4 T cells (CD45^+^ CD3^+^ CD4^+^) of total CD45^+^ cells of wild-type (WT) and ATG5^LEC-KO^ mice injected with B16F10 cells treated with 250µg isotype antibody or 250µg α-PD-1 antibody twice per week starting the moment tumors reached 100mm^3^. Mean ± SEM, each data point represents one mouse, n≥15 mice per group analyzed using Kruskal-Wallis test corrected for multiple comparisons using Dunn’s test. **E)** Frequency of intratumoral NK cells (CD45^+^ CD3^-^ NK1.1^+^) of total CD45^+^ cells of wild-type (WT) and ATG5^LEC-KO^ mice injected with B16F10 cells treated with 250µg isotype antibody or 250µg α-PD-1 antibody twice per week starting the moment tumors reached 100mm^3^. Mean ± SEM, each data point represents one mouse, n≥16 mice per group analyzed using one-way ANOVA test corrected for multiple comparisons using Kruskal-Wallis test corrected for multiple comparisons using Dunn’s test. **F)** Frequency of intratumoral KLRG1^+^ CD8 T cells of total CD8 T cells of wild-type (WT) and ATG5^LEC-KO^ mice injected with B16F10 cells treated with 250µg isotype antibody or 250µg α-PD-1 antibody twice per week starting the moment tumors reached 100mm^3^. Mean ± SEM, each data point represents one mouse, n≥13 mice per group analyzed using Kruskal-Wallis test corrected for multiple comparisons using Dunn’s test. **G)** Frequency of intratumoral IFNγ^+^ CD8 T cells of total CD8 T cells of wild-type (WT) and ATG5^LEC-KO^ mice injected with B16F10 cells treated with 250µg isotype antibody or 250µg α-PD-1 antibody twice per week starting the moment tumors reached 100mm^3^. Mean ± SEM, each data point represents one mouse, n≥11 mice per group analyzed using the Kruskal-Wallis test corrected for multiple comparisons using Dunn’s test. **H)** Gating strategies for Granzyme B^+^, Ki67^+^, KLRG1^+^, and IFNγ^+^ CD8 T cells.

**Supplementary Figure 8.**

**A)** Representative immunofluorescent images and quantification of intratumoral LYVE1 (magenta) and CD31 (green) double-positive vessels from wild-type (WT) and ATG5^LEC-KO^ mice bearing B16F10 and B16F10VEGF-C tumors. Nuclei are stained with DAPI. Mean ± SEM, n≥5 tumors from independent mice analyzed using Brown-Forsythe and Welch ANOVA test corrected for multiple comparisons using Dunnett’s T3 multiple comparisons test. Scale bars 100 µm. **B)** Tumor growth curve of individual wild-type (WT) and ATG5^LEC-KO^ mice injected with B16F10VEGF-C cells. n=8. **C)** Tumor growth curve of wild-type (WT) and ATG5^LEC-KO^ mice injected with B16F10VEGF-C tumors. Mean ± SEM, n=8 mice per group analyzed using two-way ANOVA corrected for multiple comparisons using Sidak’s test. **D)** Number of intratumoral Tregs, immature Myeloid cells (imm-Myeloid), granulocytic myeloid-derived suppressor cells (MDSCs), monocytic MDSCs, and myeloid dendritic cells (DCs) in B16F10VEGF-C tumors. Mean ± SEM, each data point represents one mouse, n≥7 analyzed using unpaired t-test with Welch’s correction or Mann-Whitney test (monocytic MDSCs). **E)** Number of intratumoral Tregs, immature Myeloid cells (imm-Myeloid), granulocytic myeloid-derived suppressor cells (MDSCs), monocytic MDSCs, and myeloid dendritic cells (DCs) in B16F10 tumors. Mean ± SEM, each data point represents one mouse, n=5 analyzed using unpaired t-test with Welch’s correction. **F)** Number of naïve (CD44^low^ CD62L^high^) CD4 and CD8 T cells in TdLNs from wild-type (WT) and ATG5^LEC-KO^ mice bearing B16F10VEGF-C tumors. Mean ± SEM, each data point represents one mouse, n≥6 analyzed using unpaired t-test (CD4) with Welch’s correction (CD8). **G)** Median fluorescence intensity (MFI) of S1PR1 on naïve CD4 and CD8 T cells in TdLNs from wild-type (WT) and ATG5^LEC-KO^ mice bearing B16F10VEGF-C tumors. Mean ± SEM, each data point represents one mouse, n≥3 analyzed using unpaired t-test (CD8) with Welch’s correction (CD4). **H)** Number of NK cells (CD3^-^ NK1.1^+^) in TdLNs from wild-type (WT) and ATG5^LEC-KO^ mice bearing B16F10VEGF-C tumors. Mean ± SEM, each data point represents one mouse, n≥7 analyzed using unpaired t-test. **I)** Tumor growth curve of individual wild-type (WT) and ATG5^LEC-KO^ mice injected with B16F10VEGF-C cells treated with 250µg isotype antibody or 250µg α-PD-1 antibody twice per week starting the moment tumors reached 100mm^3^. n=13, 11, 13, 11 respectively. **J)** Tumor growth curve of wild-type (WT) and ATG5^LEC-KO^ mice injected with B16F10 VEGF-C cells treated with 250µg isotype antibody or 250µg α-PD-1 antibody twice per week starting the moment tumors reached 100mm^3^. Mean ± SEM, n=11-13 mice per group analyzed using two-way ANOVA corrected for multiple comparisons using Sidak’s test.

**Video 1. 3D view of T cells in inguinal LN of WT mice at steady-state**

Video of a representative 3D reconstitution of a cleared whole inguinal lymph node from a wild-type (WT) mouse harboring widespread tdTomato fluorescence expression (magenta) in cells/tissues at steady-state. Lymph nodes were stained for CD3 (green). Corresponding to Fig. 1B.

**Video 2. 3D view of T cells in inguinal LN of ATG5^LEC-KO^ mice at steady-state**

Video of a representative 3D reconstitution of a cleared whole inguinal lymph node from an ATG5^LEC-KO^ mouse harboring widespread tdTomato fluorescence expression (magenta) in cells/tissues at steady-state. Lymph nodes were stained for CD3 (green). Corresponding to Fig. 1B.

**Video 3. Intravital timelapse of labeled naïve CD4 T cells in inguinal LN of WT mice at steady-state**

Representative videos of intravital imaging of inguinal lymph nodes of wild-type (WT) mice, taken at the indicated time points. Corresponding to Fig. 2D.

**Video 4. Intravital timelapse of labeled naïve CD4 T cells in inguinal LN of ATG5^LEC-KO^ mice at steady-state**

Representative videos of intravital imaging of inguinal lymph nodes of ATG5^LEC-KO^ mice, taken at the indicated time points. Corresponding to Fig. 2D.

## Method details

### Cell Culture

HDLECs were commercially purchased from Promocell (C-12217), cultured on dishes precoated with 0.1% gelatin (Sigma Aldrich), and used between passages 2 and 10. HDLECs were grown in ECGMV2 and added with SupplementMix (C-22211 and C-39226, Promocell). SiRNA transient transfection was performed twice on consecutive days using 40 nM, non-targeting siRNA (si CTRL) and siRNA against human ATG5 (si ATG5), ULK1 (si ULK1) (D-001810, L-004374, L-005049 respectively). Treatments with 25μM chloroquine (CQ) were done for 48h. B16F10 and B16F10 VEGF-C cell lines by Prof. Tatiana Petrova and HCmel12mCherry cells by Prof. Lukas Sommer and all grown in RPMI-1640 medium (R8758, Sigma Aldrich) with 10% Fetal Bovine Serum (FBS, Hyclone, Thermo Fisher Scientific). All cells were maintained in an incubator at 37 °C with 5% CO2 and 95% air.

### Immunoblot analysis

HDLECs were lysed in a modified Laemli buffer (125 mM Tris-HCl, pH 6.8 buffer containing 2% SDS and 20% glycerol) with the addition of protease and phosphatase inhibitors (A32953, A32957 respectively, Thermo Fisher Scientific). Proteins were separated using SDS-PAGE under reducing conditions, transferred to a nitrocellulose membrane, and examined by immunoblotting. Primary antibodies used were rabbit anti-ATG5 (12994S, CST), rabbit anti-GAPDH (2118S, CST), rabbit anti-ULK1 (8054S, CST), rabbit anti-SPT (ab236900, Abcam), rabbit anti-SPNS2 (ab59972, Abcam) and rabbit anti-SPHK1 (ab71700, Abcam). Appropriate secondary antibodies were from Cell Signaling Technology and Thermo Fisher Scientific (Erembodegem, Belgium). Images were acquired using the Amersham Imager 680 (*GE Healthcare* Life Sciences). Quantification of western blot data was done using Image Studio Lite software (v5.2).

### Quantitative-Real-Time PCR

RNeasy Plus mini kit (74136, Qiagen) was used for RNA extraction, and reverse transcription kit QuantiTect (205313, Qiagen) for cDNA generation. Gene expression was determined with ORA qPCR Green L mix (QPD0105, HighQu) utilizing the QuantStudio™ 5 Real-Time PCR System (Thermo Fisher Scientific) and analyzed using the delta-delta Ct method. Primer sequences are available in Table 1.

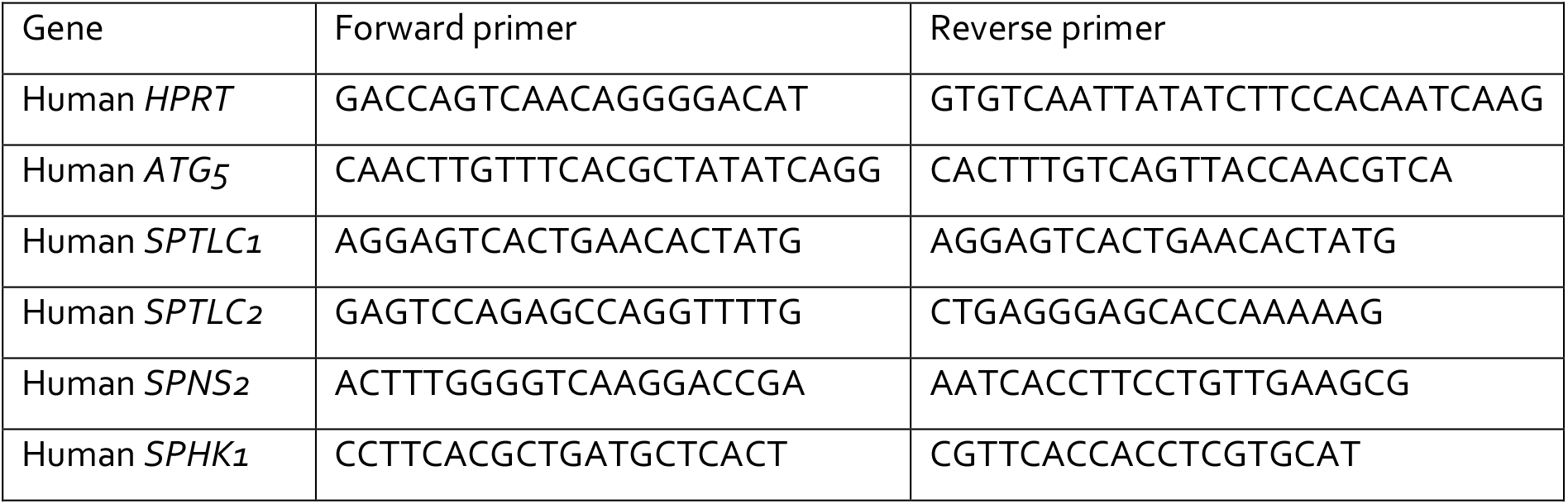

### Enzyme-linked immunosorbent assay (ELISA)

Medium was collected from si CTRL, si ATG5, si ULK1, or CQ-treated HDLECs after 8h incubation with endothelial medium containing 10%charcoal-stripped serum (A3382101, Thermo Fisher Scientific). S1P ELISA was performed following manufacturers’ protocol (MBS2700637, MyBiosource).

### Immunofluorescent staining

HDLECs were seeded on coverslips at the desired confluence. siRNA transient transfection was performed following the protocol described above. The next day, plasmid (mCherry-hLC3B-pcDNA3.1, addgene plasmid #40827) transfection (800 ng/well) was performed with Lipofectamine 2000 (Invitrogen, 16688), following manufacturer’s protocol. Cells were fixed with 4% paraformaldehyde (J19943K2, Thermo Fisher Scientific) 24h after plasmid transfection. Cells were stained overnight at 4°C with mouse anti-PDI (ab2792, Abcam) and rabbit anti-SPT (ab23696, Abcam). Cells were washed and incubated with corresponding secondary antibodies (A-21235 and A-21244 respectively, Thermo Fisher Scientific) for two hours at room temperature. Cells were rinsed with PBS, stained with DAPI for 5 min, and mounted using Prolong^®^ Gold Antifade Reagent. Images were recorded on a Zeiss LSM 780 – SP Mai Tai HP DS. Colocalization was analyzed using JACoP plugin Manders coefficients or Coloc 2 Pearson coefficient from ImageJ/Fiji Software.

### Mouse models

Animal procedures were approved by the Institutional Animal Care and Research Advisory Committee (KU Leuven) (P167/2018, P187/2018, P052/2021) and were performed in accordance with the institutional and national guidelines and regulations. Mice from the EC-specific inducible Cre-driver line *Prox1*-cre*^ERT2^* (71) were crossed with *Atg5*^fl/fl^ mice (72) to obtain mice with LEC-specific deletion of the *Atg5*gene. These lines were on a 100% C57BL/6 background. For experiments in this study, we used mice expressing Cre (Prox1-Cre+*^ERT2^*; *Atg5^fl/fl^*), referred to as ATG5^LEC-KO^ and their Cre-negative littermates (Prox1-Cre-*^ERT2^*; *Atg5^fl/fl^*), referred to as WT. Tamoxifen (T5648, Sigma Aldrich) injection (i.p.50 mg/kg) was done daily for 5 consequent days 1 week prior to experimental procedures. These lines were on a 100% C57BL/6 background. 4-deoxypyridoxine-HCl (DOP, D0501, Sigma Aldrich) treated mice received drinking water with 10g/L glucose plus 100mg/L DOP, vehicle-treated mice received drinking water with 10g/L glucose.

### Tumor cell injections

Subcutaneous tumor cell injection of 150.000 B16F10, HCMel12mCherry, or B16F10 VEGF-C cells in 100 μL PBS was performed and tumor growth was followed up several times per week. For immunotherapy experiments, 250μg of anti-PD1 or anti-B-galactosidase (clone RMP1- 14 and clone GL117 respectively, Polpharma Biologics) or 100µg of anti-CTLA4 (clone 4F10, Polpharma Biologics) or polyclonal Armenian hamster IgG (BE0091, BioXcell) was injected i.p. starting on the day tumors reached 100mm^3^ and repeated twice a week. Mice were sacrificed when tumors reached a maximum size of 1500 mm^3^.

### Integrin blocking assay

Mice were injected intravenously with 100 μg monoclonal antibody to integrin α4 (clone PS/2; BioXCell) and 100 μg monoclonal antibody to integrin αL (clone M17/4; BioXCell) and sacrificed eight hours later.

### Intravital imaging

(Prox1-Cre+*^ERT2^*; *Atg5^fl/fl^*), referred to as ATG5^LEC-KO^, and their Cre-negative littermates (Prox1-Cre-*^ERT2^*; *Atg5^fl/fl^*), referred to as WT, were crossed with R26-mTmG mice expressing the mTmG construct, leading to widespread cell membrane-localized tdTomato fluorescence expression in tissues/cells. CD4 naïve T cells were isolated from pooled lymph nodes and spleens using the naïve CD4 T cell isolation kit (130-104-453, Miltenyi Biotec) and stained with BioTracker 655 Red Cytoplasmic Membrane Dye (SCT108, Sigma Aldrich). 2 Million cells were injected intravenously into mice two hours prior to imaging. HEVs were identified by intravenous injection of 10µg anti-MECA79 antibody (53-6036-82, Invitrogen). Two hours after injection, mice were anesthetized using isoflurane inhalation (2%–23% isoflurane/oxygen mixture). Mice were placed in a supine position, while anesthesia was maintained and the right flank around the 4th mammary gland was shaved and disinfected with a piece of gauze soaked in Chlorhexidine 0.5% in ethanol (EtOH) 70%. An incision was made close to the nipple of the 4th mammary gland with lateral and diagonal directions until the inguinal lymph node was located. The inguinal lymph node was exposed by separating the skin and underlying fat pad through blunt dissection. Then, the fatty layer on top of the inguinal lymph node was carefully removed with sharp forceps until the inguinal lymph node was exposed. A gauze soaked in sterile 1x PBS was placed on top of the tissue to keep the tissue hydrated. After surgical preparation, mice were placed in a facemask within a custom-designed imaging box, on top of an inlay with a hole covered by a coverslip on the bottom. Mice were placed in such a position that the inguinal lymph node was located in the middle of the coverslip. Isoflurane was introduced through the facemask and ventilated by an outlet on the other side of the box and kept at 1-2% isoflurane/oxygen mixture for the duration of imaging. The imaging box and microscope were kept at 32 °C by a climate chamber surrounding the entire stage of the microscope including the objectives. Imaging was performed on an inverted Leica SP8 Dive system (Leica Microsystems, Mannheim, Germany) equipped with 3 tunable HyD-RLD hybrid detectors and an InSight X3 laser (SpectraPhysics) using the Leica Application Suite X (LAS X) software. All images were acquired at 12-bit with a 25×water immersion objective with a free working distance of 2.40 mm (HC FLUOTAR L 25x/0.95 W VISIR 0.17). To obtain time-lapse movies, lymph nodes were imaged every 5 minutes using a z-step of size 2.5 µm for a total duration of 2 hours. Endogenously expressed TdTomato was excited with a wavelength of 1050nm and was detected from 580nm to 620nm. GFP and BioTracker 655 Red Cytoplasmic Membrane Dye were excited with a wavelength of 780nm and were detected from 500nm to 550nm and 650 to 700nm respectively. Lateral displacement analysis was performed using ImageJ. Images of different time points (1h interval) representing the same Z-stack were merged and displacement was measured by drawing a line between the same cell in the different time points. Analysis of inflowing and emigrating cells was done using ImageJ.

### Tissue processing

Thymus, spleen, and pool of LNs (inguinal, axillary, brachial, and mesenteric) were mechanically disrupted and filtered through 40µm strainers (43-50040-51, Pluriselect) to obtain single-cell solutions. Lymph fluid was collected from cisterna chyli with a needle as previously described (73) and blood samples were collected from the submandibular vein. Tumor tissues were transferred into a gentleMACS C tube (130-093-237, Miltenyi Biotec) containing digestion medium (KnockOut™ DMEM (10829018, Thermo Fisher Scientific), penicillin/streptomycin (P4333, Sigma Aldrich), 2x Antibiotic-Antimycotic (A5955, Sigma Aldrich), 1 mM sodium pyruvate (S8636, Sigma Aldrich), 1x MEM Non-Essential Amino Acids Solution (M7145, Sigma Aldrich) supplemented with 0.1% collagenase II (17101015, Thermo Fisher Scientific), 0.25% collagenase IV (17104019, Thermo Fisher Scientific) and 15 μg/mL DNase (D4527-10KU, Sigma Aldrich) I. Each sample was further dissociated using the gentle MACS dissociator system (Miltenyi Biotec). Further purification of immune cells was done by magnetic enrichment of the CD45+ fraction (130-052-301, Miltenyi Biotec). All cell suspensions were kept in PEB buffer (0.5% BSA and 2 mM EDTA in PBS).

### Flow cytometry

Cells were incubated with FC block (101319, Biolegend) for 10 minutes and then stained with Fixable Viability Dye eFluor™ 780 (65-0865-14, Thermo Fisher Scientific), anti-CD3 (130-116-530, Miltenyi Biotec), anti-CD4 (130-116-546, Miltenyi Biotec), anti-CD8 (565968, BD Biosciences), anti-CD44 (130-116-495, Miltenyi Biotec), anti-CD62L (565261, BD Biosciences), anti-S1PR1 (FAB7089P, R&D systems), anti-CD19 (115545, Biolegend), anti-NK1.1 (746876, BD Biosciences), anti-CD45 (748370, BD Biosciences), anti-CD25 (740714, BD Biosciences), anti-Ki67 (652424, Biolegend), anti-CD69 (741478, BD Biosciences), anti-KLRG1 (138423, Biolegend), anti-GranzymeB (130-116-486, Miltenyi Biotec) following the FoxP3/Transcription factor staining buffer set (00-5523-00, Thermo Fisher Scientific). To assess apoptotic cells, cells were stained with propidium iodide (PI) (P4170, Sigma). Cells were analyzed using a BD FACSSymphony A5 within 24h of staining and data was analyzed using FlowJo software (v10.8.1).

### Cell sorting

At least 9 TdLNs per condition were pooled and digested following the protocol described in (74). Cells were incubated with FC block for 30 minutes and stained with anti-MECA79 (53-6036-82, Invitrogen), anti-GP38 (127423, Biolegend), anti-CD31 (551262, Biolegend), anti-CD45 (103116, Biolegend) and anti-Ter119 (116208, Biolegend) for one hour on ice. Cells were washed and 7AAD (00-6993-50, Invitrogen) was added just prior to sorting. Endothelial cells were identified as single 7AAD^-^, Ter119^-^, CD45^-^, CD31^+^ cells and immune cells as single 7AAD^-^, Ter119^-^, CD45^+^ cells.

### BD Rhapsody™ Single-Cell Analysis System

Both samples were loaded on a BD Rhapsody cartridge with a targeted capture of 6000 cells. Reverse transcription, cDNA amplification, and library construction were performed following the manufacturer’s instructions (23-24117(02)). The final pooled libraries were sequenced on a NovaSeq6000 flow cell (Illumina), spiked with 20% PhiX control DNA to increase the sequence complexity.

### Single-cell transcriptomic analysis

The BD Rhapsody Sequence Analysis Pipeline (BD; version 1.11) was used to map the FASTQ files to the mouse reference genome (mm10-2020-A). Single-cell analysis was performed using RStudio (R version 4.0.5) starting from the complete, unfiltered matrix according to the workflow proposed by the Marioni and Theis lab (75). Cells with less than 200 genes expressed and genes expressed in less than 3 cells were filtered out of the count matrix. Outliers were identified based on 3 metrics: number of expressed genes, library size and mitochondrial proportion. Cells 5 MADs (Median Absolute Deviation) away from the median value were filtered out for the number of expressed genes and library size, and 15 MADs were used as an upper limit for the mitochondrial proportion, resulting in lenient filtering. After a first clustering round, extra low-quality cells were removed based on matrix origin, gene expression, and library size. The samples were merged using the Seurat merge function, counts were normalized and log2 transformed using the NormalizeData function, both from the Seurat R package (v4.0.2) using default parameters. Detecting highly variable genes, scaling, finding clusters, and creating UMAP plots was done using the Seurat pipeline. Clustering was performed using the first 20 principal components and a resolution of 0.8. Differential gene expression analysis between cell clusters and conditions was performed using the Wilcoxon rank-sum test. Intracellular signaling was predicted using NicheNet (51). NicheNet is a computational method using prior knowledge of ligand-to-target signaling paths to predict ligand-receptor interactions in the cells of interest. To carry out transcription factor network inference, analysis was performed as described (47) using the SCENIC R package (version 1.1.0).

### Immunohistochemistry

Mouse LN tissue samples were immediately fixed in 4% PFA overnight at 4°C, dehydrated, and embedded in paraffin. Immunostainings were performed using an in-house protocol. Antibodies used were goat anti-LYVE1 (R&D systems, AF2125), rabbit anti-CD3 (ab5690, Abcam), rat anti-PNAd (120802, Biolegend), rat anti-CD31 (550274, BD), and the signal was enhanced using TSA Plus CY3/CY5 detection kits (NEL744001KT and NEL705A001KT, Akoya Biosciences). Images were acquired using an inverted microscope (IX73, Olympus). Analyses were performed with *Olympus cellSens Dimension* software. For other stainings, mouse LN tissue samples were fixed in 2% PFA overnight and embedded in a 4% agarose solution in PBS. Tissues were stained with rabbit anti-ATG5 (12994S, CST), rabbit anti-LC3B (3868S, CST), goat anti-LYVE1 (R&D systems, AF2125) and rabbit anti-ACKR4 (PA5-33408, Thermo Fisher Scientific) overnight at 4°C and corresponding secondary antibodies were added for 3 hours at 4°C. Confocal images were acquired with a Leica sp8x confocal microscope (KUL-VIB CCB), 60X magnification.

### MILAN staining

Multiple Interactive Labeling by Antibody Neodeposition (MILAN) immunohistochemistry was performed according to a previously published method (76,77). Tissue sections (3µm) were prepared from FFPE murine lymph node samples. Following dewaxing, antigen retrieval was performed using PT link (Agilent) using 10 mM EDTA in Tris-buffer pH 8. Immunofluorescence staining was performed using Bond RX Fully Automated Research Stainer (Leica Biosystems) with the following primary antibodies: anti-F4/80 (ab6640, Abcam), anti-CD3 (ab16669, Abcam), anti-CD4 (# 14-9766-82, Invitrogen), anti-CD31 (ab28364, Abcam), anti-MECA79 (53-6036-82, Thermo Fisher), anti-LYVE1 (R&D systems, AF2125), anti-VEGFR3 (# 14-5988-82, Thermo Fisher), and anti-FoxP3 (# 14-5773-82).The sections were incubated for 4 hours with the primary antibodies, washed, and then incubated for 30 min with secondary antibodies. A coverslip was placed into the slides with a medium containing 4,6-diamidino-2-phenylindole (DAPI) and scanned using a Zeiss AxioScan Z.1 (Zeiss) at 10X magnification. The coverslips were removed after 30 minutes soaking in washing buffer. Antibody stripping was performed in a buffer containing 1% SDS and β- mercaptoethanol for 30 minutes at 56°C. The staining procedure was repeated for several rounds until all markers were stained and scanned.

### MILAN image analysis

Image analysis was conducted using a previously described (78). Briefly, images were corrected for the field-of-view artifact using a method described in the literature (79), Then, the overlapping regions of adjacent tiles were stitched together by minimizing the Frobenius distance. Next, images from consecutive rounds were aligned (registered) following an algorithm previously described (80). To register the images from consecutive rounds, the first round was set as a fixed image while all the following rounds were used as moving images. The DAPI channel was used to calculate transformation matrices, which were then applied to the other channels. The quality of the overlapping regions was visually evaluated and poor registered areas were removed from downstream analyses. Following registration, a region of interest was selected for the next parts of the analysis. Then, tissue autofluorescence was removed by subtracting a baseline image with only a secondary antibody. Finally, cell segmentation was applied to the DAPI channel using a finetuned model for STARDIST (81) which delineates a contour for each cell present in the tissue. For each of these cells, the following features were extracted: topological features (X/Y coordinates), morphological features (nuclear size), and molecular features (Mean Fluorescence Intensity (MFI) of each measured marker).

### MILAN single cell analysis

Mean Fluorescence Intensity (MFI) values were normalized using Z-scores within each sample, as recommended in Caicedo et al. (82). To avoid a strong influence from outliers in downstream analyses, Z-scores were trimmed within the [-5, 5] range. Three different clustering methods were used to map single cells to known cell phenotypes: PhenoGraph (83), FlowSom (84), and KMeans, which were implemented in the Rphenograph, FlowSOM, and stats R packages, respectively. While FlowSom and KMeans required the number of clusters as input, PhenoGraph could be executed by exclusively defining the number of nearest neighbors to calculate the Jaccard coefficient 20 (standard value). The number of clusters identified by PhenoGraph was then used as an argument for FlowSom and KMeans. Clustering was performed exclusively on a subset of the identified cells (10,000), which were selected by stratified proportional random sampling and using the following markers: CD3, CD31, CD4, CD8, FOXP3, F480, LYVE1, CD19, CD49, and MECA79. For each clustering method, clusters were mapped to known cell phenotypes based on manual annotation by domain experts. For every cell, if two or more clustering methods agreed on the assigned phenotype, the cell was annotated accordingly.

Following consensus clustering, 4 different cell types were identified (+Blank): High Endothelial Venules (MECA79+ | CD31+), CD4 T cells (CD4+ | CD3+), CD8 T cells (CD8+ | CD3+), Lymphatic Vessels (LYVE1+), Regulatory T cells (FOXP3+ | CD4+), Macrophages (F480+), Blood Vessels (CD31+), B cells (CD19+) and Natural Killer cells (CD49+). To extrapolate the cell labels to the remaining cells in the dataset, a uMap was built by sampling 500 cells for each identified cell type in the consensus clustering, and the entire dataset was projected into the uMap using the base predict R function. For each cell, the label of the closest 100 neighbors was evaluated in the uMap space, and the most frequent cell type was assigned as the label. Digital reconstructions of the tissue samples were obtained by coloring the segmentation mask with the assigned cell label. These reconstructions were further used to annotate different areas of interest: B cell follicles, T cell zone, and medulla.

Relative cytometry enrichment was performed for each area using Wilcoxon rank-sum tests. Statistical analysis and data presentation were performed using R Studio (version RStudio 2022.07.2).

### Lymph node clearing and staining

Inguinal lymph nodes were dissected and optically cleared using the FLASH tissue clearing protocol as previously described (85). Lymph node optical clearing process was performed for 3 days in total. For immunostaining, whole-cleared tissues were blocked for 1hour and incubated with a primary anti-CD3 rabbit antibody (ab5690, Abcam) for 3 days at room temperature. Subsequently, whole-cleared tissues were washed for 3-4h-hours and incubated with a secondary Alexa Fluor™ 647 donkey anti-rabbit antibody (A-31573, Invitrogen) for another 3 days in room temperature. All incubation steps were performed on a rocking plate.

### 3D imaging of lymph nodes

Optically cleared inguinal lymph nodes were placed in between two coverslips, and imaged on an upright Leica SP8 confocal microscope. Endogenously expressed TdTomato was excited at 555nm and collected at 580 – 630nm, and Alexa647 was excited at 635 nm and collected at 650–700 nm. The inguinal lymph nodes were imaged in 3D using a large-scale tile scan with a total Z-stack of approximately 200 µm with a Z-step size of 3 µm.

### Quantification and statistical analysis

All data are represented as mean ± SEM. The normality of data was checked using Anderson-Darling, D’Agostino & Pearson, and Shapiro-Wilk testing. Statistical significance between the two groups was determined by standard unpaired t-test with F-testing or one sample t-test. Unless otherwise indicated, statistical significance between multiple groups was determined by one-way ANOVA to ensure comparable variance, then individual comparisons were performed by Dunn’s posthoc test. In the case of non-normality, Mann-Whitney or Kruskal Wallis tests were performed. Analysis was done in Prism v9.0f, GraphPad. GP p value style was used. * represents a *p*-value < 0.05, ** < 0.01, *** *p* < 0.001 and **** p < 0.0001 where a *p*-value < 0.05 is considered significant.

## Acknowledgments

We are thankful to the VIB Single Cell Core, VIB Flow Core Leuven, and VIB Nucleomics for support and access to the instrument park (vib.be/core-facilities). We thank N.D. Lakic, P. Nazari, J. Lamote, and J. Verhoeven for excellent technical support. We thank T. Petrova and and B. Prat Luri for the generating and providing us the B16F10 VEGF-C cell line. We thank Taija Mäkinen for the *Prox1-CreERT2* mice and Noboru Mizushima for Atg5^fl/fl^ mice. Images were recorded on a Zeiss LSM 780 – SP Mai Tai HP DS (Cell and Tissue Imaging Cluster (CIC), Supported by Hercules AKUL/11/37 and FWO G.0929.15 to Pieter Vanden Berghe, University of Leuven. P.A. is supported by grants from the Flemish Research Foundation (FWO-Vlaanderen; G076617N, G049817N, G070115N), the EOS DECODE consortium N° 30837538, the EOS MetaNiche consortium N° 40007532, Stichting tegen Kanker (FAF-F/2018/1252) and the iBOF/21/053 ATLANTIS consortium with G.B. D.H. is the recipient of an FWO Doctoral Fellowship from the Flemish Research Foundation (FWO-Vlaanderen, 1155121N), Belgium. K.J. is the recipient of an FWO Postdoctoral Fellowship from the Flemish Research Foundation (FWO-Vlaanderen, 12Y4322N).

