## Supplementary Figures for "An autophagy program that promotes T cell egress from the lymph node controls responses to immune checkpoint blockade"

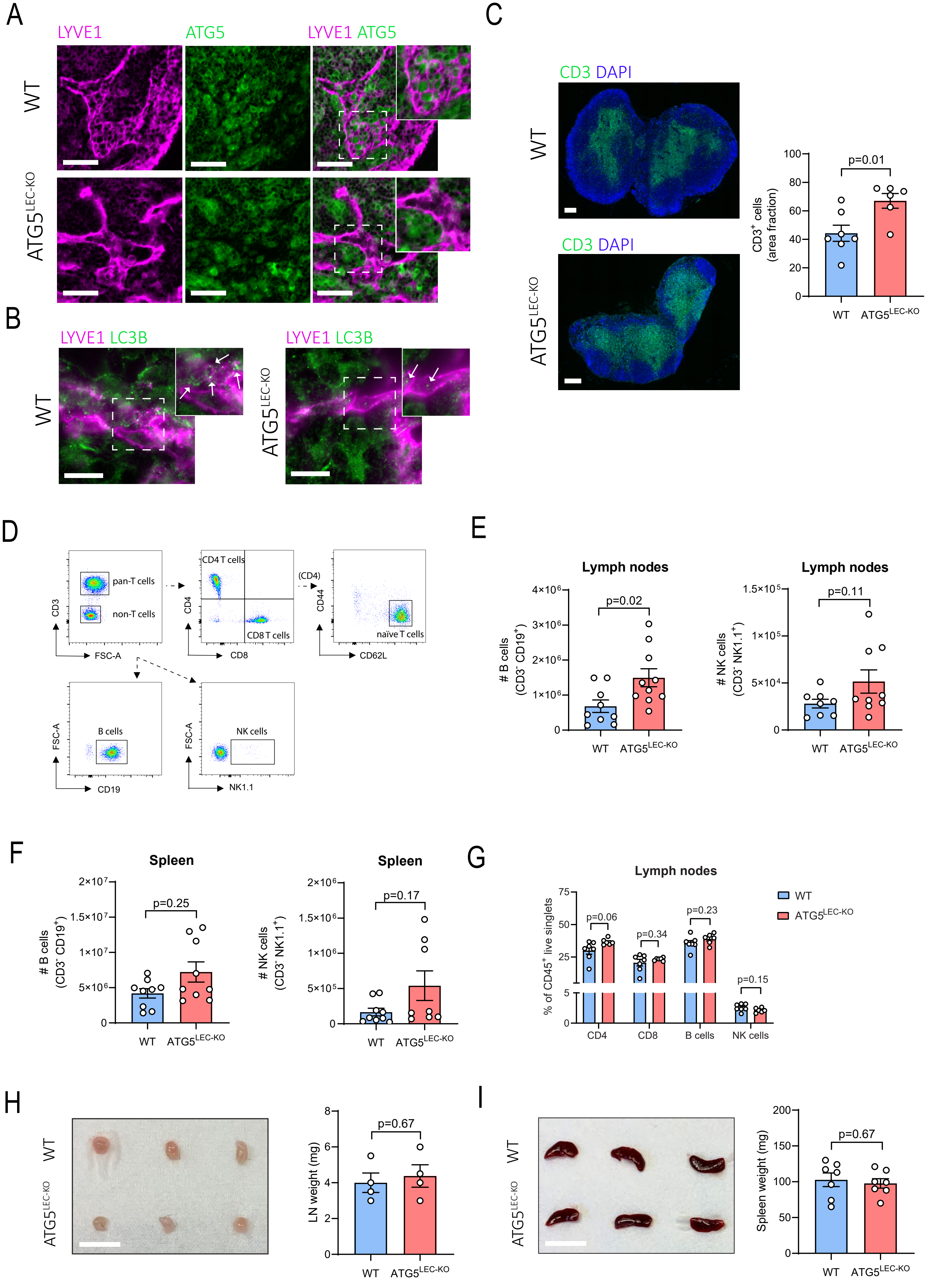


Supplementary Figure 1

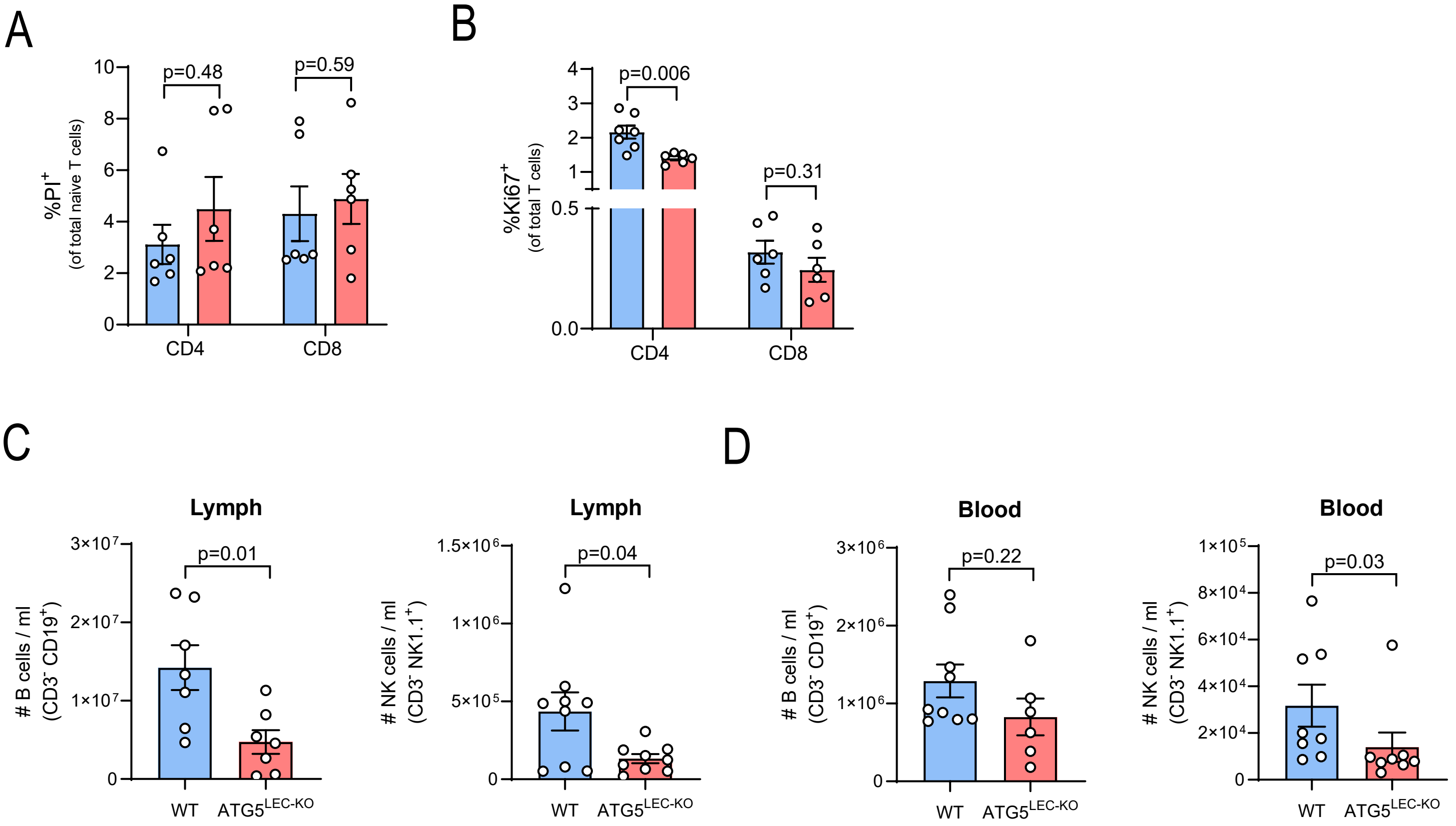


Supplementary Figure 2

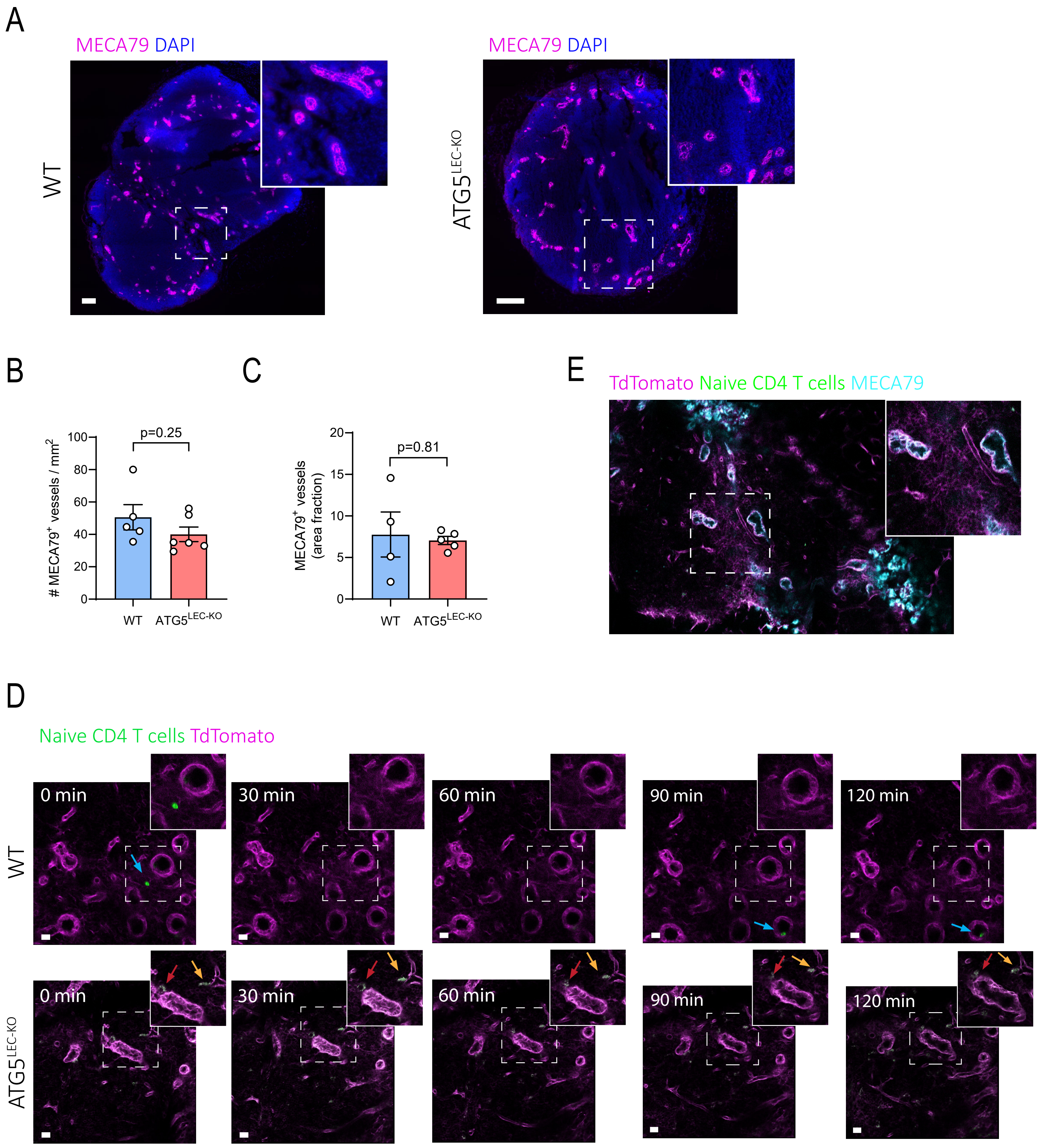


Supplementary Figure 3

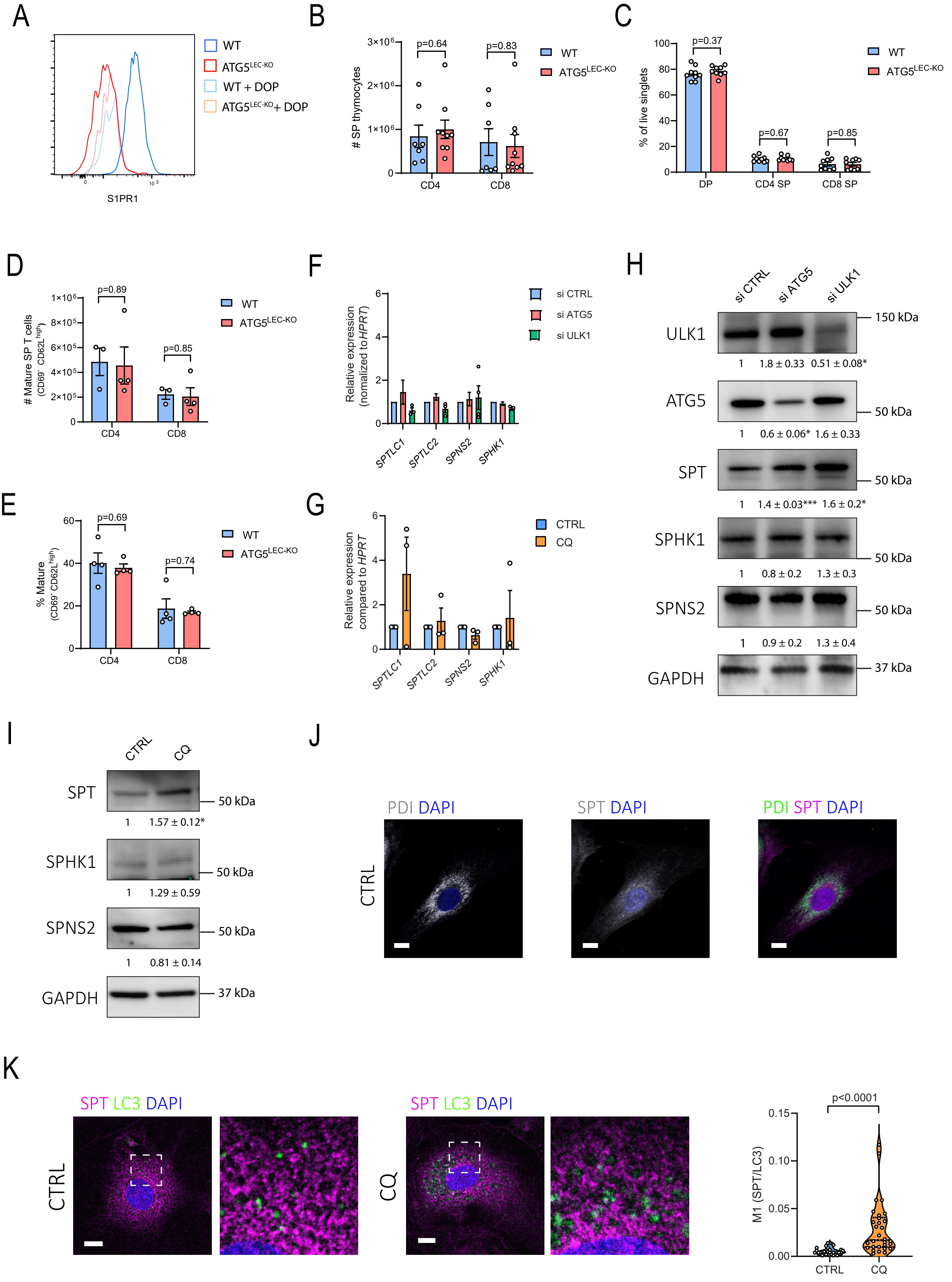


Supplementary Figure 4

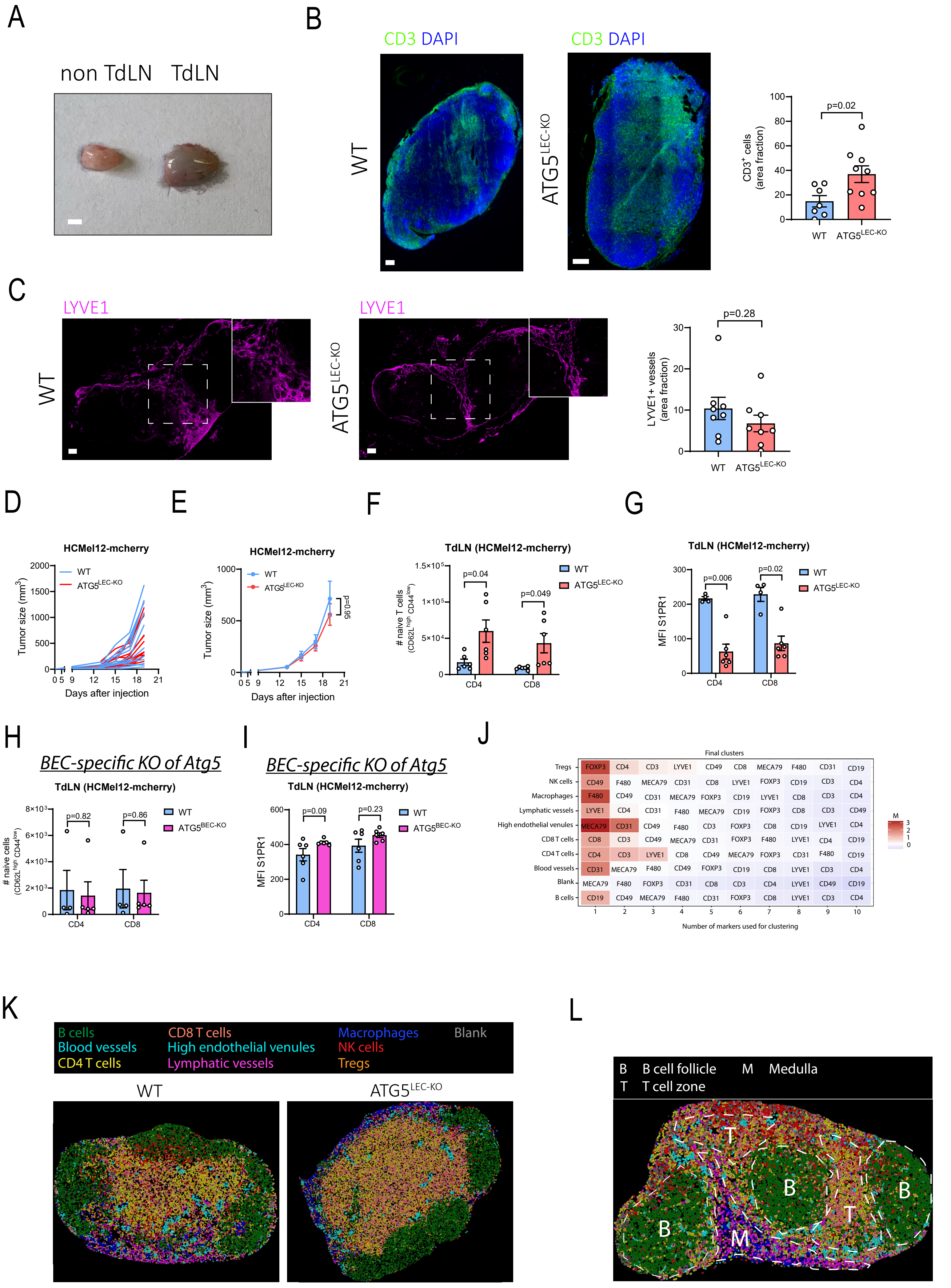


Supplementary Figure 5

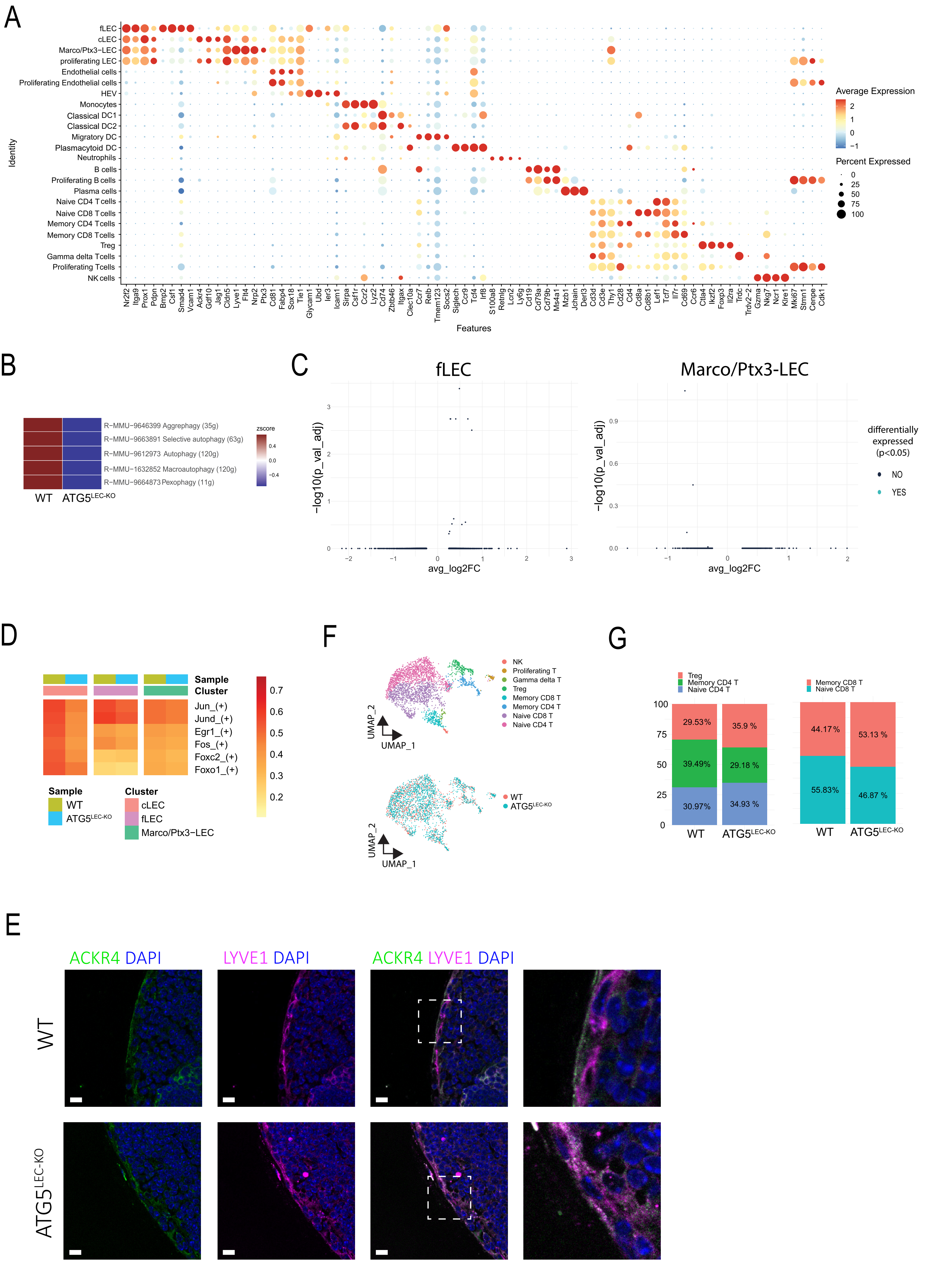


Supplementary Figure 6

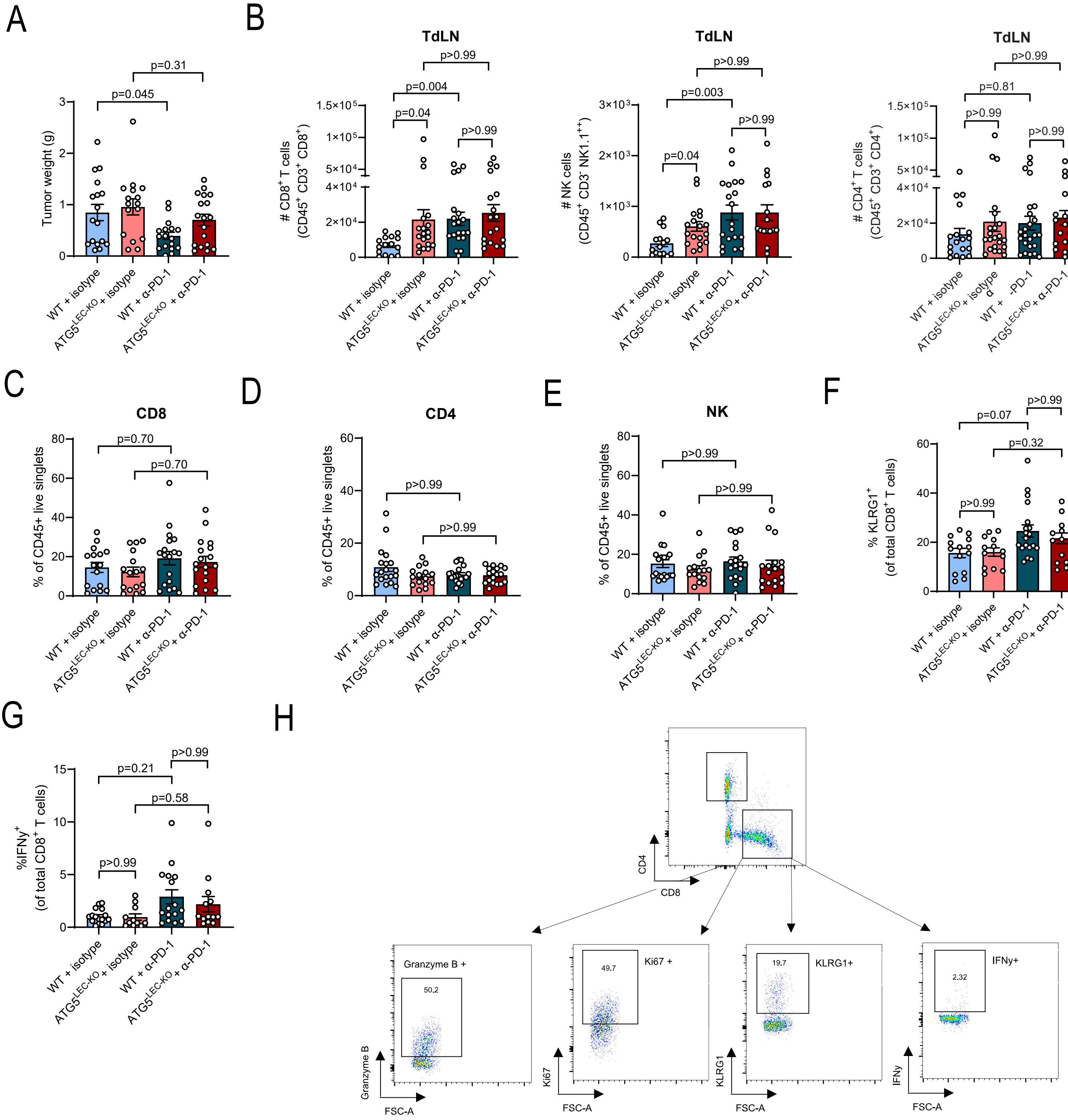


Supplementary Figure 7

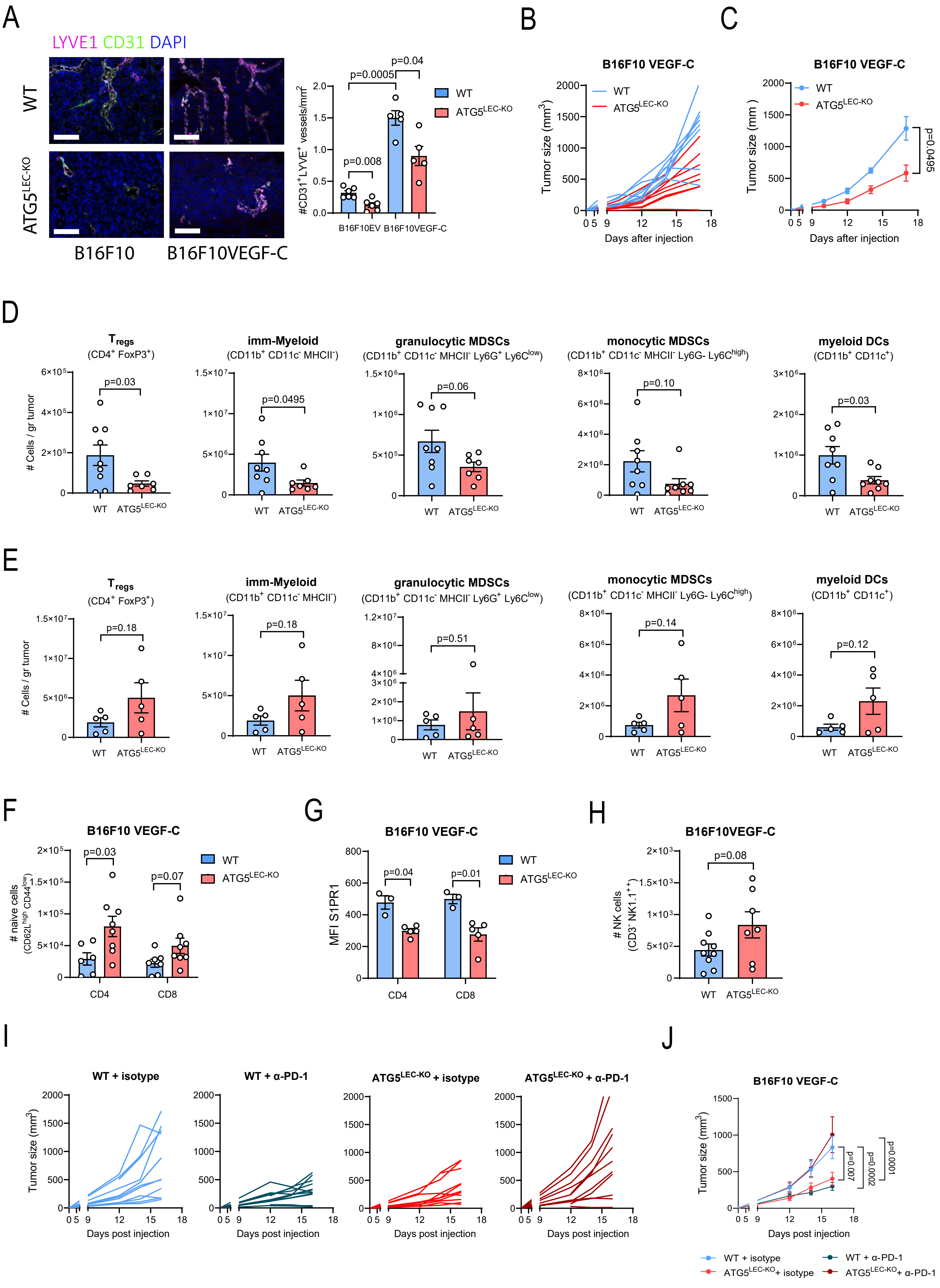


Supplementary Figure 8
